# Human FACT complex coordinates cohesin and transcription on chromatin

**DOI:** 10.64898/2026.10.06.754922

**Authors:** Jonay Garcia-Luis, Andres Canela, Valeriya Malysheva, Toyoaki Natsume, Mohammad M Karimi, Masato T. Kanemaki, Mikhail Spivakov, Luis Aragon

**Affiliations:** DNA Motors Research Group, MRC Laboratory of Medical Sciences (LMS), Du Cane Road, London W12 0NN, United Kingdom; Institute of Clinical Sciences, Imperial College, London W12 0NN, United Kingdom; Radiation Biology Centre. Kyoto University, Yoshida-Konoe-Cho, Sakyo-Ku, Kyoto 606-8501, Japan; Functional Gene Control Group. MRC London Institute of Medical Sciences (LMS), Du Cane Road, London W12 0NN, United Kingdom; Department of Chromosome Science, National Institute of Genetics, Research Organization of Information and Systems (ROIS), Mishima, Shizuoka, 411-8540, Japan; Graduate Institute for Advanced Studies, SOKENDAI, Mishima, Shizuoka 411-8540, Japan; Department of Biological Science, Graduate School of Science, The University of Tokyo, Bunkyo-ku, Tokyo 113-0033, Japan; Departmento de Ciencias Médicas Básicas and Instituto de Tecnologías Biomédicas, Universidad de La Laguna, La Laguna, Spain; Computational Neurobiology lab, VIB Center for Molecular Neurology, VIB, Antwerp, Belgium; Department of Biomedical, Pharmaceutical and Veterinary sciences, University of Antwerp, Antwerp, Belgium; Research Center for Genome & Medical Sciences, Tokyo Metropolitan Institute of Medical Science, Setagaya-ku, Tokyo 156-8506, Japan; Comprehensive Cancer Centre, School of Cancer & Pharmaceutical Sciences, Faculty of Life Sciences & Medicine, King’s College London, London, United Kingdon

## Abstract

The spatial organisation of eukaryotic genomes is critical for coordinating DNA-dependent processes such as transcription and replication. Cohesin contributes to this organisation by extruding chromatin loops; however, how cohesin achieves processive translocation along nucleosome-dense chromatin in vivo remains poorly understood. Here, we identify the histone chaperone FACT as a regulator of cohesin dynamics in human cells. We show that the FACT subunit SUPT16H colocalises and interacts with cohesin on chromatin. Depletion of FACT impairs cohesin translocation from its loading sites, leading to reduced chromatin looping and decreased boundary strength of topologically associating domains (TADs), as revealed by Hi-C. In addition, FACT depletion results in the accumulation of cohesin within gene bodies that are not engaged in loop formation, which is associated with reduced transcription. Importantly, degradation of cohesin in FACT-depleted cells restores transcriptional output, indicating that cohesin mislocalisation underlies this effect. Together, these findings demonstrate that FACT facilitates cohesin translocation along chromatin, promoting efficient loop extrusion while preventing its aberrant accumulation within transcribed regions. This work provides mechanistic insight into how chromatin structure modulates cohesin function to coordinate genome organisation with transcription in mammalian cells.

## Introduction

Eukaryotic genomes are packaged into chromatin, in which DNA is wrapped around histone octamers to form nucleosomes, creating a compact yet dynamic structure that both protects and restricts access to the genetic material. To regulate DNA accessibility, cells deploy chromatin remodelling complexes and histone chaperones that modulate nucleosome organisation and composition. These factors can reposition nucleosomes along DNA, evict histones, or exchange histone variants, thereby facilitating access to DNA for transcription and other essential DNA-dependent processes (Rippe et al., 2007).

FACT (Facilitates Chromatin Transcription) is a heterodimeric histone chaperone consisting of Spt16 (SUPT16H in humans) and Pob3 (SSRP1 in humans) (Orphanides et al., 1998) that allows RNA Polymerase II (Pol II)-driven transcription elongation through DNA templates containing nucleosomes *in vitro* (Orphanides et al., 1998). FACT interacts with H2A-H2B dimers and stabilizes partially disassembled nucleosomes generated during transcription (Hsieh et al., 2013; Liu et al., 2020). By facilitating nucleosome reorganization ahead of the transcribing polymerase and promoting nucleosome reassembly behind it, FACT helps Pol II traverse nucleosome-bound DNA while preserving chromatin integrity (Ehara et al., 2022; Hsieh et al., 2013; Tsunaka et al., 2016). Consistent with its role in transcription, genome-wide analyses in various organisms have shown that FACT is enriched at actively transcribed genes *in vivo* (Chang et al., 2018; Saunders et al., 2003; True et al., 2016). Surprisingly, knockdown of FACT in human cells does not affect global transcription, instead, only a subset of genes exhibit significant up or down-regulation (Li et al., 2007; Mylonas and Tessarz, 2018; Wang et al., 2024). The role of FACT in facilitating RNA Pol II transcription in cells, and why a subset of genes are differentially affected remains unclear.

In yeast cells, FACT also supports cohesin function by facilitating its translocation along chromatin (Garcia-Luis et al., 2019). Cohesin is a multimeric complex involved in the organisation of chromatin during interphase as well as in the establishment and maintenance of sister chromatid cohesion from S phase to mitosis (Yatskevich et al., 2019). Cohesin is a tetrameric ring complex formed by the SMC ATPases Smc1 and Smc3, the kleisin subunit Scc1 (RAD21 in mammalian cells), and a fourth subunit Scc3 (STAG1/2 in mammalian cells)(Losada et al., 1998).. Cohesin’s role in genome organisation depends on its ability to form large chromatin loops (Rao et al., 2017) through an activity termed loop extrusion (Davidson et al., 2019; Kim et al., 2019).

Cohesin loading onto chromosomes requires the Scc2/4 loader complex (Ciosk et al., 2000; Watrin et al., 2006). Importantly, Scc2 also stimulates the ATPase activity of cohesin (Petela et al., 2018), and it is necessary for loop extrusion *in vitro* (Davidson et al., 2019; Kim et al., 2019). In vertebrate genomes, cohesin-mediated loops are preferentially terminated at convergent binding sites for the insulator protein CTCF, leading to the demarcation of topologically associating domains (TADs). The N-terminal domain of CTCF can physically interact with the STAG1/2-RAD21 subcomplex of cohesin, serving as a barrier that halts loop extrusion at CTCF sites (Li et al., 2020; Nora et al., 2020; Parelho et al., 2008). Consistently, depletion of cohesin (Rao et al., 2017) or the cohesin loading factor NIPBL causes TADs and loops to disappear globally (Haarhuis et al., 2017), confirming that NIPBL is required for ongoing loop extrusion *in vivo*. In contrast, degradation of another cohesin regulator, WAPL, leads to an increase in loop size (Haarhuis et al., 2017).

NIPBL ChIP signals are enriched at promoters and enhancers of active genes, suggesting that cohesin can be recruited to transcription start sites through interactions between the cohesin loader and the Mediator transcriptional complex (Kagey et al., 2010). However, studies in yeast indicate that cohesin loading occurs throughout the bodies of actively transcribed genes (Petela et al., 2018). More recently, observations from yeast and mammalian systems suggest that elongating RNA polymerase II can act as a moving barrier to loop extrusion, pushing or repositioning cohesin as it transcribes along genes and thereby contributing to its enrichment at promoters. (Banigan et al., 2022; Jeppsson et al., 2022). Once loaded, cohesin is expected to engage in active loop formation until it reaches CTCF sites in convergent orientation. As cohesin loop extrudes through chromatin, it needs to overcome DNA-bound obstacles, including nucleosomes and other active protein complexes such as transcribing RNA polymerases. *In vitro*, human cohesin has been reported to bypass obstacles larger than its ring size 50% of the time while extruding loops (Pradhan et al., 2022); however, surprisingly, it is blocked by individual nucleosomes when diffusing on DNA (Davidson et al., 2016). In yeast and human cells, cohesin has been reported to be enriched between convergently oriented genes (Busslinger et al., 2017; Lengronne et al., 2004), which has been proposed to reflect that active RNA polymerase II both pushes cohesin to these sites and represents an obstacle for cohesin movement when encountered in the opposite direction (Banigan et al., 2022; Jeppsson et al., 2022). Besides transcription, the replication machinery also constrains cohesin translocation on chromosomes (Dequeker et al., 2022).

Chromatin remodellers have also been shown to facilitate both the loading and function of cohesin in chromatin. The RSC (Remodeling the Structure of Chromatin) complex interacts with the Scc2/4 loader complex and generates a nucleosome-free region onto which cohesin can load (Muñoz et al., 2019). Another remodeller important for cohesin function is the FACT complex (Garcia-Luis et al., 2019). *In vivo* experiments in yeast have shown that inactivation of FACT disrupts cohesin-dependent TADs (Garcia-Luis et al., 2019). Given FACT’s roles in transcription elongation (Orphanides et al., 1998) it has been proposed that cohesin translocation (or loop extrusion) requires FACT-mediated passage through nucleosomes (Garcia-Luis et al., 2019).

An open question is how FACT affects cohesin movement and loop extrusion on chromatin and whether this is an evolutionary conserved function. Here, we tested whether FACT facilitates cohesin function in human cells and investigated the underlying mechanisms. Cohesin and FACT co-localise in a large proportion of sites. Importantly, depletion of the FACT subunit SUPT16H causes a genome-wide decrease in cohesin binding peaks, particularly those away from its loading sites, as well as a reduction in the number of loops and the strength of TADs. Surprisingly, SUPT16H depletion also causes the accumulation of cohesin within gene-bodies of transcribed genes. The presence of cohesin on these genes correlates with a decrease in transcription levels that is associated with its accumulation along the gene-bodies, demonstrating that RNA polymerase II cannot push cohesin when it accumulates on DNA. These results highlight an intricate relation between cohesin, FACT, and transcription, that needs to be coordinated.

## Results

### Cohesin and FACT co-localise across the genome

Yeast FACT destabilises and partially unfolds nucleosomes (Liu et al., 2020; Valieva et al., 2016) and its inactivation leads to increased nucleosome occupancy at transcription start sites (TSS) (True et al., 2016). In mouse embryonic stem cells, FACT is similarly enriched at TSS (Mylonas and Tessarz, 2018). Consistent with this role, rapid FACT depletion in human cells disrupts nucleosome integrity within transcribed regions and impairs transcription through chromatin, supporting a model in which FACT in facilitates RNA polymerase II passage while preserving nucleosome structure (Žumer et al., 2024). Notably, the cohesion loader NIPBL is also enriched at promoter regions (Kagey et al., 2010), suggesting potential spatial coordination between FACT and cohesion.

To investigate this, we analysed the genome-wide distribution of human FACT (hFACT) alongside NIPBL and the cohesin subunit RAD21 in human HCT116 cells. Using calibrated ChIP-seq, we measured the occupancy of the hFACT subunit SUPT16H across gene bodies and compared these with transcriptional activity measured by nascent RNA sequencing. SUPT16H occupancy positively correlates with nascent transcription levels (Fig.1 a, b) and displays a characteristic metagene profile marked by strong enrichment at the TSS and a sustained signal across gene bodies (Fig.1 c). We next assessed co-enrichment between FACT and cohesin components, NIPBL, RAD21 and CTCF (Fig.1 d). A substantial fraction (∼67%) of SUPT16H peaks overlap with RAD21 peaks (Fig.1d, e; Fig.S1) indicating extensive co-localisation. Together, these data reveal a strong genome-wide correlation between FACT occupancy and cohesin binding sites in human cells.

**Figure 1.**
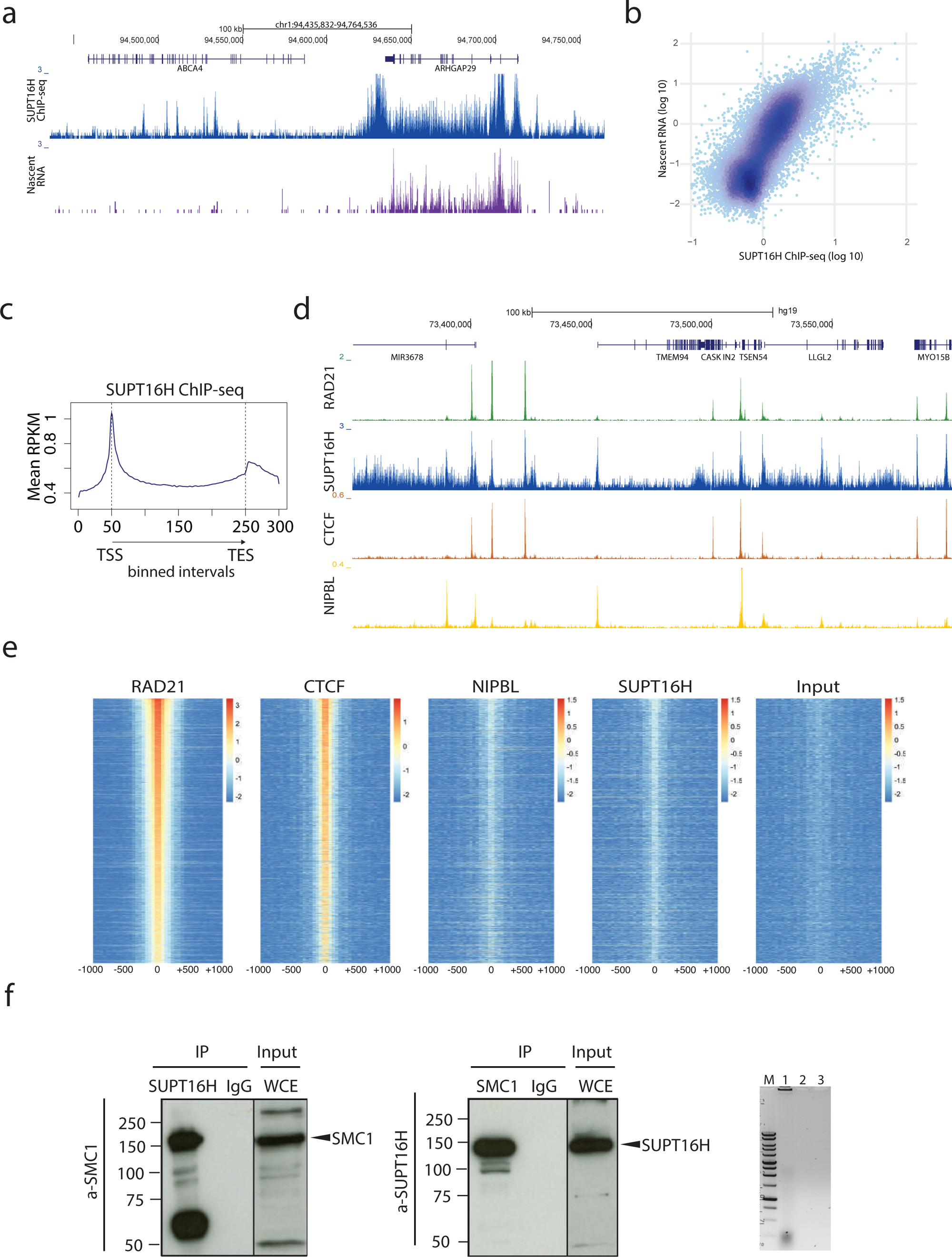
Genome-wide co-localization and interaction of FACT and cohesin in human cells. **a**, Calibrated ChIP-seq profiles of SUPT16H (FACT) and nascent RNA levels across an active genomic locus in HCT116 cells. Gene annotations are shown above. SUPT16H signal correlates with transcriptional activity and is enriched across gene bodies. **b**, Scatter plot showing genome-wide correlation between SUPT16H ChIP-seq signal at gene bodies and nascent RNA levels, indicating that FACT occupancy correlates with transcriptional activity. **c**, Metagene analysis of SUPT16H ChIP-seq signal across annotated genes. Gene bodies (transcription start sites (TSS) to transcription end sites (TES)) were divided in 200 equal-length bins, and flanking regions upstream of the TSS and downstream TES in 50 intervals each; genes shorter than 2 kb were excluded. SUPT16H is enriched at TSSs and has a sustained signal across gene bodies. **d**, Calibrated ChIP-seq genomic profiles of RAD21 (cohesin), SUPT16H (FACT), CTCF and NIPBL at a representative genomic locus. Overlapping binding patterns indicate that FACT and cohesin occupy common chromatin regions, including sites associated with CTCF and the cohesin loader NIPBL. **e**, Heatmaps of RAD21, CTCF, NIPBL, SUPT16H binding centred on RAD21 peaks summits (±1 kb), sorted by RAD21 signal intensity. SUPT16H co-enriches at a large fraction of cohesin-bound sites, supporting widespread co-localization between FACT and cohesin complexes. **f**, Co-immunoprecipitation analysis of SUPT16H and SMC1 from whole-cell extracts. Immunoprecipitation (IP) with antibodies against SUPT16H (left) or SMC1 (centre) followed by immunoblotting against SMC1 or SUPT16H respectively, shows reciprocal co-purification of SMC1 and SUPT16H. Cell extracts were treated with benzonase prior to IP to eliminate DNA-mediated interactions (right). IgG controls and whole-cell extracts (WCE) are shown.

A physical interaction between Spt16 and Smc1 has previously been reported in yeast (Garcia-Luis et al., 2019). Given the observed co-localisation of SUPT16H and RAD21 (Fig.1 e; Fig.S1), we tested whether this interaction is conserved in human cells. We performed reciprocal co-immunoprecipitation experiments from exponentially growing mammalian cells using antibodies against SMC1 or SUPT16H, followed by immunoblotting of the precipitates. Cell extracts were treated with benzonase to eliminate DNA-mediated interactions prior to immunoprecipitation. Immunoprecipitation of SMC1 resulted in co-purification of SUPT16H, and conversely, immunoprecipitation of SUPT16H co-purified SMC1 (Fig.1 f).

These results demonstrate that FACT and cohesin not only co-localise across the genome but also physically interact in mammalian cells, supporting a conserved functional relationship between these complexes.

### Depletion of FACT compromises cohesin translocation

Previous studies assessing the impact of FACT on transcription have relied on siRNA-mediated depletion of the SUPT16H subunit (Li et al., 2007). To directly examine the role of FACT in cohesin localisation, we instead an auxin-inducible degron (AID) system to achieve rapid and acute degradation of SUPT16H in human cells (Natsume et al., 2016). In this system, the plant derived ubiquitin ligase TIR1 is constitutively expressed under the CMV promoter, enabling auxin-dependent polyubiquitination and proteosomal degradation of AID-tagged proteins (Natsume et al., 2016).

We generated HCT116 colorectal carcinoma cells expressing SUPT16H fused to an AID tag and mClover (SUPT16H-AID-mClover) (Fig.2 a). To suppress leaky degradation arising from TIR1, we added TIR1 inhibitor auxinole to the control cells (Yesbolatova et al., 2019). Efficient degradation of SUPT16H following auxin addition was confirmed by fluorescence microscopy, western blotting and calibrated ChIP-Seq, with substantial depletion observed within 30 minutes (Fig.2 b,c; Fig.S2 a,b). First, we assessed whether loss of FACT affects the overall level of chromatin-bound cohesin using chromatin fractionation assays. Notably, the total amount of cohesin associated with chromatin remained unchanged upon SUPT16H degradation (Fig.2 c), indicating that FACT is dispensable for cohesin loading onto chromosomes. We next examined whether FACT influences cohesin distribution by performing ChIP-seq for the RAD21 cohesin subunit before and after auxin treatment. Depletion of SUPT16H resulted in a reduction in the intensity of a subset of RAD21 peaks (Fig.2 d; Fig.S2 c), suggesting that although cohesin loading is unaffected, its positioning along chromatin is altered in the absence of FACT. To investigate this further, we analysed RAD21 occupancy at CTCF-bound sites and across the intervening regions. As cohesin accumulates at CTCF sites following loop extrusion, these positions provide a proxy for cohesin translocation efficiency. In control conditions, RAD21 was strongly enriched at CTCF sites, with relatively low signal across intergenic regions (Fig.2 e). Upon SUPT16H degradation, RAD21 enrichment at CTCF sites was markedly reduced, accompanied by a modest but consistent increase in signal between these sites, as quantified by mean RPM and Auxin/Auxinole ratios (Fig.2 e). Consistent with this, analysis of RAD21 signal emanating from putative loading sites (RAD21⁺, NIPBL⁺, CTCF⁻) revealed a progressive reduction in RAD21 enrichment at increasing genomic distances in SUPT16H-depleted cells relative to controls (Fig.S2 d). Together, these findings demonstrate that while FACT is not required for cohesin loading, it is critical for efficient cohesin translocation along chromatin, thereby shaping its genome-wide distribution.

**Figure 2.**
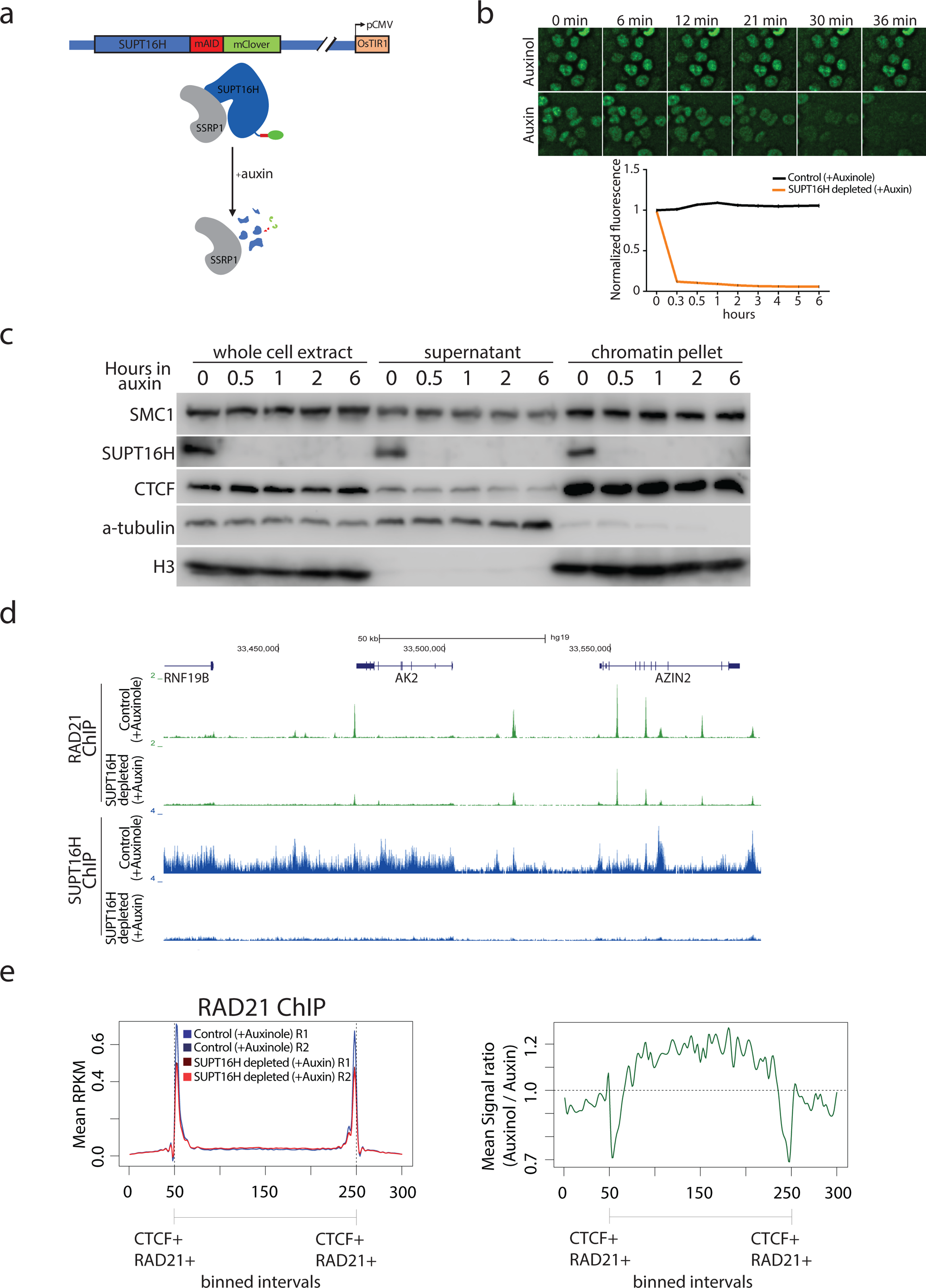
FACT depletion impairs cohesin translocation without affecting chromatin loading. **a**, Schematic of the auxin-inducible degron (AID) system used to acutely deplete the FACT subunit SUPT16H in HCT116 cells. SUPT16H was tagged with an AID degron fused to mClover, and OsTIR1 was constitutively expressed from the CMV promoter. Upon auxin addition, SUPT16H is rapidly ubiquitinated and degraded, disrupting the FACT complex. **b**, Time-course fluorescence microscopy of SUPT16H-mAID-mClover signal following auxin addition, compared to control (+auxinole). Representative images are shown at the indicated time points. Quantification of normalized fluorescence intensity, measured by live-cell imaging using an Incucyte system, demonstrates rapid depletion of SUPT16H within ∼30 min of auxin treatment. **c**, Chromatin fractionation followed by immunoblotting of whole-cell extract. Soluble (supernatant), and chromatin-bound (chromatin pellet) fractions at the indicated times after auxin addition are shown. SMC1 levels in the chromatin fraction remain largely unchanged upon SUPT16H depletion. SUPT16H is efficiently depleted, while CTCF remains chromatin-associated. α-tubulin and histone H3 serve as fractionation controls. **d**, Calibrated ChIP-seq genomic profiles of RAD21 and SUPT16H in control (+auxinole) and SUPT16H depleted (+auxin) cells at a representative locus. Upon SUPT16H depletion, RAD21 peak intensity is reduced at discrete sites, consistent with altered cohesin distribution. **e**, **Left**, aggregate analysis of RAD21 ChIP-seq signal across intervals defined by the outermost coordinates of pairs of CTCF-bound RAD21 peaks (CTCF+ RAD21+) flanking a putative cohesin-loading site (CTCF− RAD21+ NIPBL+), before (blue) and after addition of Auxin (red). **Right**, ratio of RAD21 signal (Auxin/Auxinole) showing reduced enrichment at CTCF sites and a relative increase in signal across the intervening regions after SUPT16H depletion.

### FACT facilitates the formation of chromatin loops and TADs

Cohesin is loaded onto DNA at NIPBL-bound sites and extrudes loops bidirectionally until it encounters convergent CTCF sites that define TAD boundaries (Fudenberg et al., 2016; Nora et al., 2017). Consistently, depletion of the cohesin subunit RAD21 abolishes loop formation (Rao et al., 2017).

Given that our data indicate that loss of FACT subunit SUPT16H impairs cohesion translocation along chromatin (Fig.2 e; Fig.S2 d), we hypothesised that FACT inactivation would perturb higher-order genome organisation. To test this, we performed high-resolution Hi-C in HCT116 cells expressing SUPT16H-AID-mClover following auxin-induced degradation of SUPT16H. We built high resolution Hi-C libraries from cells depleted in SUPT16H and compared them with cells where SUPT16H had not been degraded. At the megabase scale, Hi-C maps revealed the expected segregation of chromatin into A and B compartments, corresponding to transcriptionally active and inactive regions, respectively (Lieberman-Aiden et al., 2009). Eigenvector decomposition showed highly similar compartment profiles between control (+auxinole) and SUPT16H-depleted (auxin-treated) cells (Fig.3 a). Consistently, saddle plot analysis demonstrated comparable enrichment of intra-compartment (A-A, B-B) and inter-compartment (A-B) interactions under both conditions (Fig.3 b), indicating that global compartmentalisation is largely unaffected by FACT depletion.

**Figure 3.**
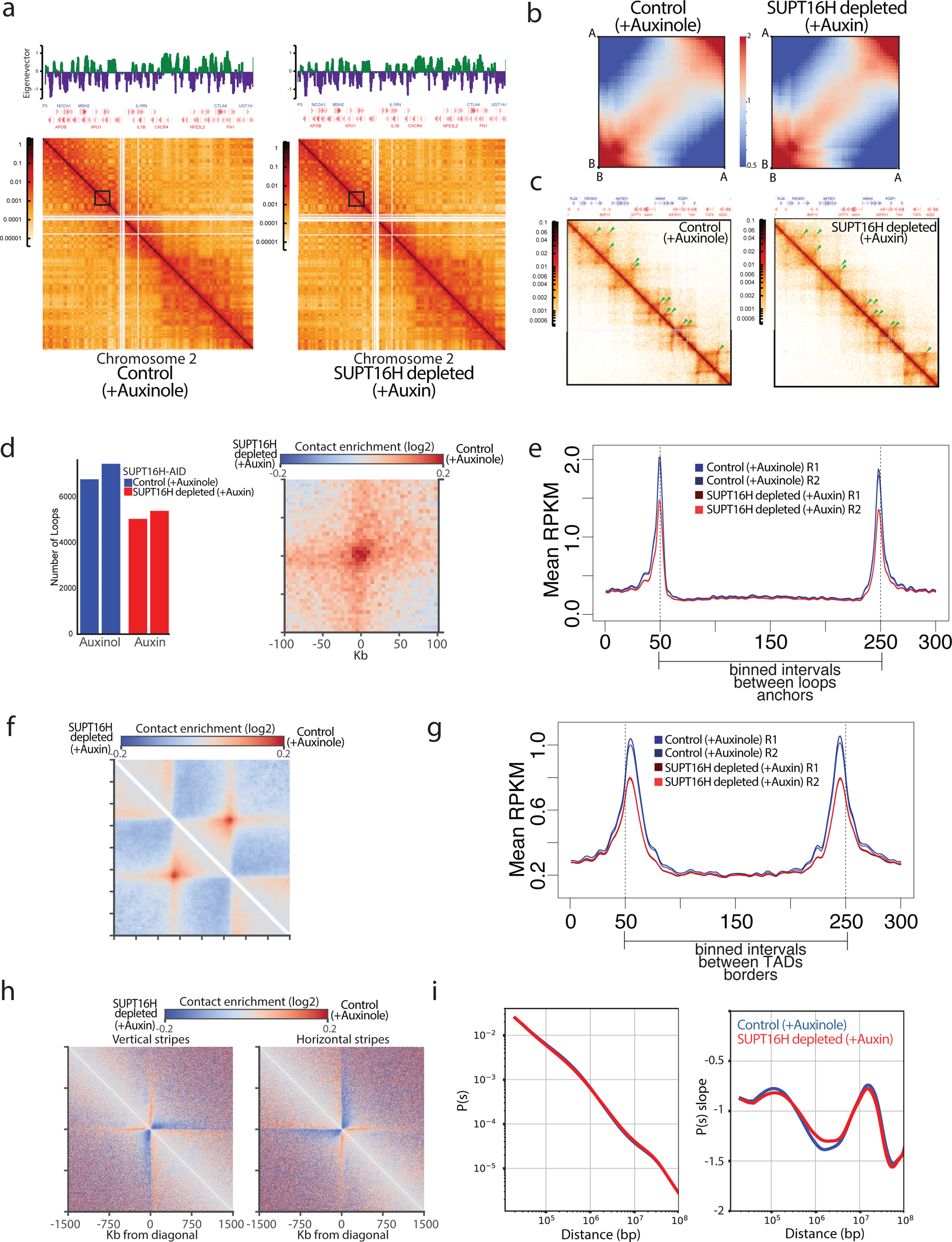
FACT promotes cohesin-dependent chromatin looping and TAD organization without altering compartment structure. **a**, Hi-C contact maps of chromosome 2 in SUPT16H-AID cells under control (+auxinole) and SUPT16H depletion (+auxin) conditions, with corresponding eigenvector (PC1) tracks indicating A/B compartment organization. Compartment profiles are largely preserved upon SUPT16H depletion. **b**, Saddle plots showing compartmentalization strength (A-A and B-B enrichment and A-B depletion) in control and auxin-treated cells. Comparable interaction patterns indicate that compartment identity and strength are not substantially affected by SUPT16H depletion. **c**, High-resolution Hi-C contact maps at a representative genomic region (inset from the boxed region in Fig.3 a) in control and auxin-treated cells. Focal interaction peaks corresponding to loops are reduced upon SUPT16H depletion. **d**, Left, number of chromatin loops identified genome-wide by HOMER in control and auxin-treated conditions. Right, aggregate peak analysis (APA) of loops showing reduced contact enrichment in auxin-treated cells. **e,** Calibrated RAD21 ChIP-seq signal across loop anchors, aggregate profiles were generated using loop outermost coordinates expanded 10kb for loops common to the control (auxinole) samples. The distance between the two loop anchors was divided into 200 equal-sized bins, with 50 flanking bins on each side. RAD21 enrichment at loop anchors is reduced upon SUPT16H depletion. **f**, Aggregate TAD analysis centred on domain boundaries, showing decreased boundary-associated contact enrichment following SUPT16H depletion. **g**, Calibrated RAD21 ChIP-seq profiles centred on TAD boundaries defined using the TADs outermost coordinates expanded 20kb for TADs common to the control (auxinole) samples. Regions were scaled as in panel e. Cohesin enrichment at TAD boundaries is reduced in SUPT16H depleted (+auxin) cells. **h**, Aggregate analysis of vertical and horizontal stripe features detected in Hi-C maps, showing reduced stripe-associated contact enrichment upon SUPT16H depletion. **i**, Genome-wide contact probability as a function of genomic distance (P(s), left) and corresponding slope (right) in control and auxin-treated cells. Profiles are largely overlapping, indicating that the characteristic genomic distance scale of chromatin folding is preserved despite local architectural changes.

In contrast, inspection of Hi-C contact maps at higher resolution revealed pronounced alterations in local chromatin architecture. While overall genome organisation was preserved, focal interaction peaks corresponding to chromatin loops were visibly reduced upon SUPT16H depletion (Fig.3 c), suggesting weakened cohesin-mediated looping. Loop calling using HOMER identified ∼7,100 loops in control cells, which decreased to ∼5,200 following SUPT16H degradation (Fig.3 d). Aggregate loop analysis confirmed a reduction in contact enrichment at loop anchors in auxin-treated cells (Fig.3 d; Fig.S3 a,b). Consistent with these observations, calibrated RAD21 ChIP-seq revealed decreased cohesin occupancy at loop anchors upon SUPT16H depletion (Fig.3 e), indicating that weakened loop signals reflect reduced cohesin accumulation at these sites.

We next examined higher-order domain organisation. TADs identified using HOMER showed reduced boundary strength following SUPT16H depletion, as evidenced by aggregate TAD analysis centred on domain boundaries (Fig.3f; Fig.S3c). This reduction in insulation was accompanied by decreased RAD21 enrichment at TAD boundaries (Fig.3g), further linking impaired cohesin positioning to altered domain architecture.

To assess the impact on cohesin processivity, we analysed architectural stripes, linear tracks of elevated contact frequency that emanate from loop anchors and reflect ongoing loop extrusion (Vian et al., 2018). Visual inspection of Hi-C maps suggested that stripe features were attenuated following SUPT16H depletion. Quantitative analysis using Stripenn (Yoon et al., 2022) confirmed a significant reduction in stripe-associated contact enrichment and overall stripe scores in auxin-treated cells (Fig. 3h; Fig. S3d), consistent with impaired cohesin translocation.

Despite these local architectural defects, the global scaling of chromatin contacts remained largely unchanged. Contact probability as a function of genomic distance (P(s)) was nearly identical between control and SUPT16H-depleted cells across all distances examined (Fig. 3i; Fig. S3e), with no detectable shift in the characteristic shoulder of the curve. This indicates that the average length scale of loop extrusion is preserved. Such behaviour is consistent with perturbations that affect cohesin translocation or barrier interactions without altering intrinsic extrusion capacity, as observed upon depletion of CTCF or chromatin-associated obstacles such as MCM complexes (Nora et al., 2017; Wutz et al., 2017).

Together, these results demonstrate that FACT is required for the proper formation of cohesin-dependent chromatin architecture. Depletion of SUPT16H weakens loop anchors, reduces loop numbers, and diminishes TAD boundary insulation, all of which correlate with decreased cohesin accumulation at these sites. Architectural stripes are similarly attenuated, indicating reduced cohesin processivity. Notably, the preservation of the P(s) profile suggests that FACT does not alter the intrinsic length scale of loop extrusion, but rather facilitates efficient cohesin translocation along chromatin, thereby promoting the formation and stabilisation of loops and TAD boundaries.

### Cohesin accumulates at transcribed genes upon FACT depletion

Cohesin is a central organiser of 3D genome architecture. Following loading by the NIPBL-MAU2 complex, cohesin redistributes from its initial deposition sites to accumulate at convergent CTCF sites, active transcription start sites (TSSs), and intergenic regions between convergently transcribed genes. This redistribution reflects a dynamic balance between WAPL-mediated release, stalling at CTCF boundaries, and transcription-driven translocation of cohesin along chromatin (Busslinger et al., 2017; Lengronne et al., 2004). RNA polymerase II (RNAPII) acts as a moving barrier to loop extrusion with head-on encounters between RNAPII and cohesin impeding cohesin progression, thereby converting actively transcribed genes into dynamic architectural boundaries (Banigan et al., 2022; Jeppsson et al., 2022).

Given the coupling between transcription and cohesin redistribution, and the established role of FACT in facilitating RNAPII elongation, we asked whether FACT depletion alters cohesin positioning within gene bodies. To address this, we performed calibrated ChIP-seq for RAD21 and the cohesin loader NIPBL in control and SUPT16H-depleted HCT116 cells. Under control conditions, cohesin displayed the expected enrichment at TSSs, with comparatively low occupancy across gene bodies (Fig. 4a,b). In contrast, depletion of SUPT16H resulted in a pronounced accumulation of RAD21 across gene bodies, as demonstrated by both representative loci and metagene analysis, with a similar trend observed for NIPBL (Fig. 4a). This redistribution affected approximately half of all genes for RAD21, whereas NIPBL exhibited a similar but more restricted pattern (Fig. 4c,d). Notably, cohesin enrichment was broad and diffuse along gene bodies rather than confined to discrete peaks, explaining why it was not captured in peak-based analyses.

**Figure 4.**
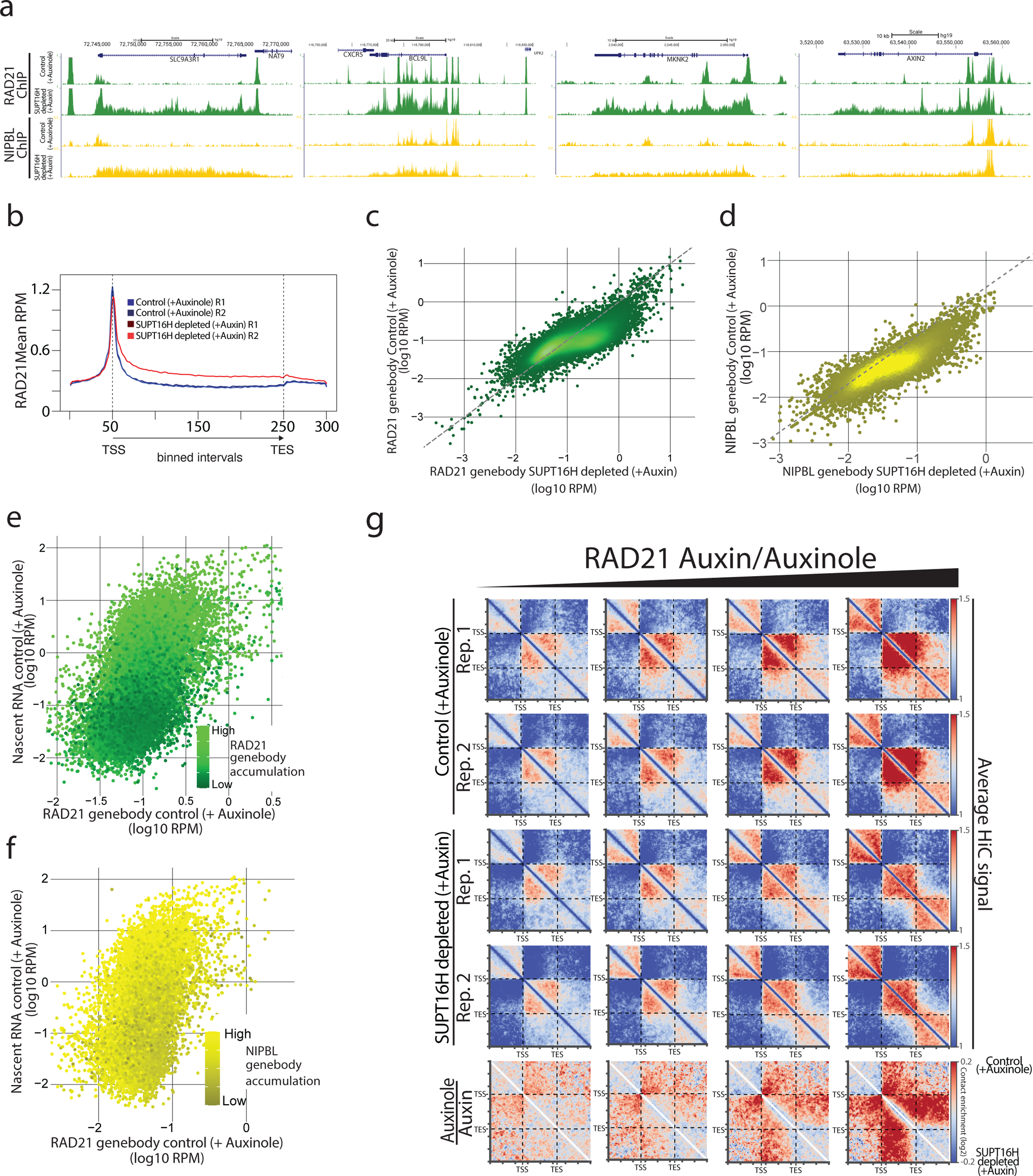
FACT depletion leads to cohesin accumulation within gene bodies and alters intra-genic chromatin interactions. **a**, Calibrated ChIP-seq of RAD21 and NIPBL in control (+auxinole) and SUPT16H depletion (+auxin) conditions at representative loci. Upon SUPT16H depletion, both RAD21 and NIPBL display increased and broadened signal across gene bodies. **b**, Metagene analysis of RAD21 ChIP-seq signal across gene bodies. The distance between transcription start site (TSS) to transcription end site (TES) was divided in 200 bins, with 50 bins in each flanking region. genes shorter than 2,000 bp were excluded. RAD21 shows increased enrichment across gene bodies following SUPT16H depletion across biological replicates. **c**, Scatter plot comparing RAD21 signal across gene bodies in control and auxin-treated conditions, indicating increased RAD21 enrichment at a subset of genes upon SUPT16H depletion. **d**, As in c, for NIPBL, showing a similar but more restricted increase in gene body enrichment following SUPT16H depletion. **e**, Scatter plot showing the relationship between RAD21 accumulation in gene bodies upon SUPT16H depletion (colour scale from dark (low accumulation) to light green (high accumulation)), nascent RNA levels prior to auxin addition, and RAD21 levels prior to auxin addition. RAD21 enrichment positively correlates with transcriptional activity. **f**, As in e, for NIPBL, showing a similar positive correlation between gene body accumulation and transcription levels. **g**, Aggregate Hi-C contact maps scaled to gene bodies (aligned from TSS to TES) stratified by RAD21 accumulation levels in control and auxin-treated cells. Genes with higher RAD21 accumulation display reduced intra-genic contact enrichment upon SUPT16H depletion.

Because cohesin dynamics within genes are tightly linked to transcription, we next examined the relationship between cohesin accumulation and transcriptional activity. RAD21 and NIPBL enrichment across gene bodies showed a strong positive correlation with nascent RNA levels, with highly transcribed genes displaying the greatest accumulation (Fig. 4e,f. Fig. S4a,b). These results indicate that FACT depletion promotes transcription-dependent retention of cohesin within gene bodies.

Chromatin interactions within and between genes are shaped by the interplay between loop-extruding cohesin and RNAPII (Banigan et al., 2022; Jeppsson et al., 2022; Rowley and Corces, 2018). Transcriptionally active genes typically exhibit increased intra-genic contacts and insulation, supported by cohesin enrichment. We therefore asked whether the accumulation of cohesin following FACT depletion enhances local chromatin folding. High-resolution Hi-C analysis centred on TSSs and transcription end sites (TESs) revealed the opposite effect. Stratifying genes by RAD21 accumulation, we observed that those with the highest cohesin enrichment exhibited the strongest reduction in intra-genic contact frequency (Fig. 4d).

To disentangle the contributions of transcription, we further stratified genes based on both RAD21 accumulation and transcriptional changes following SUPT16H depletion (Fig. S4c). Notably, the reduction in intra-genic contacts and loss of insulation persisted even at genes where transcriptional output remained unchanged, despite substantial cohesin accumulation. Extending this analysis to the domain scale, we found that although RAD21 occupancy at TAD boundaries was globally reduced upon FACT depletion (Fig. 3g), cohesin accumulation within TADs increased with transcriptional activity (Fig. S4d), indicating that transcription-dependent cohesin retention occurs across multiple levels of genome organisation.

Together, these findings support a model in which FACT facilitates cohesin translocation along chromatin. In the absence of FACT, cohesin becomes trapped within gene bodies, accumulating in a state that is impaired in loop extrusion. This leads to reduced intra-genic contacts, diminished insulation, and a global weakening of loop and stripe features, consistent with defective cohesin dynamics following SUPT16H depletion.

### Cohesin accumulation at gene bodies blocks transcription

DNA-bound proteins can act as physical barriers to elongating RNA polymerase II (RNAPII), leading to transcriptional pausing or arrest. This phenomenon is well established across systems, where transcription factors, nucleosomes, and other chromatin-associated complexes hinder polymerase progression (Candelli et al., 2018; Deuschle et al., 1990; Kireeva et al., 2002; Shukla et al., 2011). Conversely, RNAPII itself acts as a moving barrier that constrains cohesin translocation during loop extrusion (Banigan et al., 2022; Jeppsson et al., 2022), highlighting a bidirectional interplay between transcription and cohesin dynamics.

The aberrant accumulation of cohesin within gene bodies upon FACT depletion therefore raises the possibility that cohesin, in turn, interferes with transcription elongation. Because both cohesin and elongating RNAPII occupy and traverse the same chromatin template, their co-existence likely requires tight coordination. Disruption of this balance may increase collision frequency or steric interference, thereby compromising RNAPII processivity and transcriptional output.

To test whether cohesin retention feeds back onto transcription, we quantified changes in nascent RNA following SUPT16H depletion in relation to RAD21 accumulation across gene bodies (Fig. 5a). We observed a strong association between cohesin enrichment and transcriptional repression: genes with higher transcriptional activity prior to depletion exhibited greater RAD21 accumulation and underwent more pronounced reductions in transcription (Fig. 5a).

**Figure 5.**
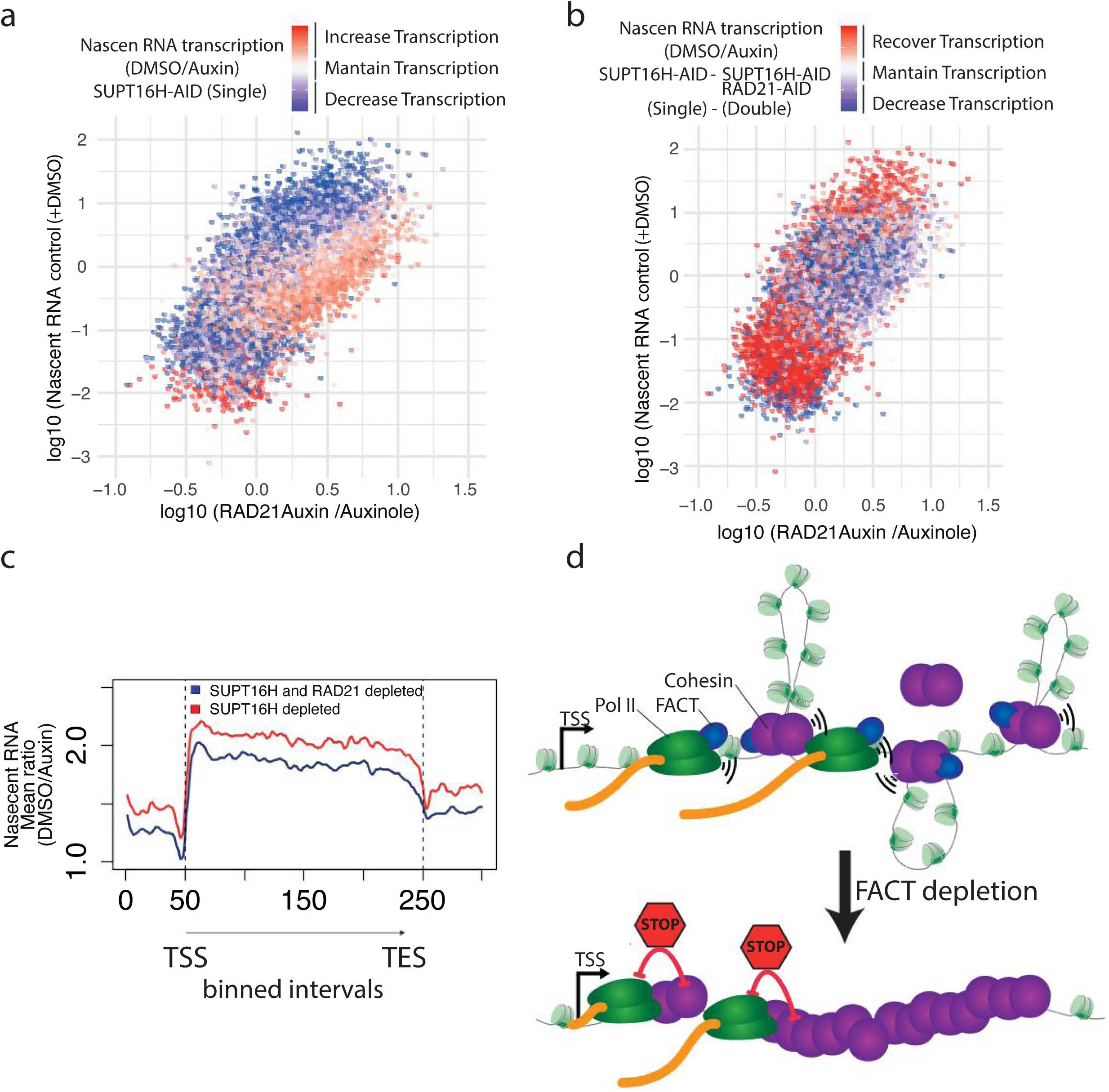
Cohesin accumulation within gene bodies impairs transcription and is rescued by RAD21 depletion. **a**, Scatter plot showing the transcriptional response to SUPT16H depletion across all expressed genes. Each gene is colour-coded by its DMSO/Auxin nascent RNA ratio in the SUPT16H-AID single degron. Higher DMSO/auxin ratios indicate larger decrease in transcription following auxin treatment, and lower ratios indicate an increase. Colours range from red (transcription increase upon depletion) to blue (transcription decreased upon depletion). Highly transcribed genes show greater RAD21 accumulation in gene bodies following SUPT16H depletion and correspondingly more pronounced transcriptional decrease, suggesting that cohesin accumulation contributes to transcriptional repression at these loci. **b,** Scatter plot comparing the transcriptional response to auxin-induced depletion between SUPT16H-AID single degron and SUPT16H-AID RAD21-AID double degron cells. Each gene is colour-coded according to the difference between DMSO/Auxin nascent RNA ratio in the single and double degrons (single - double). Positive values (in red) indicate lower DMSO/Auxin ratios in the double degron therefore a recovery of transcription upon RAD21 co-depletion, in contrast negative values (in blue) indicate a greater transcriptional decrease in the double degron. Highly transcribed genes that accumulated cohesin in gene bodies upon SUPT16H depletion show positive values (red) indicating a preferential recovery of transcription when RAD21 is simultaneously depleted, consistent with cohesin-mediated repression being causally linked to the transcriptional decrease observed upon SUPT16H loss. **c,** Metagene analysis of the DMSO/Auxin nascent RNA signal ratio across gene bodies for top 5% transcribed genes displaying cohesin accumulation upon SUPT16H depletion (RAD21 Auxin/Auxinole ratio >1). Gene bodies (TSS to TES) were divided in 200 equal size bins, with 50 bins upstream the TSS and downstream the TES. Lower DMSO/Auxin ratios indicate a smaller transcriptional decrease following auxin treatment. The SUPT16H-AID RAD21-AID double degron shows lower DMSO/Auxin ratios indicating a recovery of transcription at high transcribed genes following RAD21 co-depletion. **d**, Model illustrating the differential roles of FACT in coordinating transcription and cohesin dynamics on chromatin. In control conditions, FACT performs two complementary functions. First, it facilitates RNA polymerase II (Pol II) elongation by destabilizing and reorganizing nucleosomes, thereby promoting efficient transcription through chromatin. Second, FACT supports cohesin translocation by reducing the nucleosomal barrier to loop extrusion, enabling cohesin to move processively along chromatin after loading. In this context, elongating Pol II acts as a directional, moving barrier that redistributes cohesin along gene bodies, while FACT ensures that both Pol II and cohesin can traverse nucleosome-dense regions. This coordinated activity allows cohesin to efficiently relocate away from gene bodies and accumulate at CTCF-bound sites, supporting loop formation and normal transcription. Upon FACT depletion, these two functions are differentially affected. Impaired nucleosome reorganization reduces Pol II processivity, limiting its ability to transcribe through chromatin and to displace cohesin. In parallel, cohesin translocation becomes inefficient due to increased chromatin resistance, leading to its retention within gene bodies. As a result, cohesin accumulates in regions of active transcription where it is no longer efficiently repositioned. In this state, cohesin acts as a static obstacle to elongating Pol II, increasing steric interference and leading to transcriptional attenuation.

To directly assess causality, we generated RAD21-AID SUPT16H-AID double degron cells (Fig. S5a) and compared nascent transcription following auxin-induced co-depletion. Among genes with RAD21 accumulation upon SUPT16H depletion, the magnitude of transcriptional recovery in the double degron followed a clear gradient of transcriptional activity; most pronounced in the top 5% of transcribed genes and progressively diminishing through the top 10%, 25%, 50% and 75% percentiles (Fig. 5b,c, Fig. S5b). This gradual relationship between transcriptional activity and the degree of recovery upon RAD21 co-depletion supports a direct causal link between cohesin accumulation and transcriptional repression at actively transcribed loci.

Together, these results reveal a reciprocal functional relationship between transcription and cohesin. While elongating RNAPII redistributes cohesin along gene bodies, excessive cohesin accumulation in turn impairs transcription, likely by imposing physical or topological constraints on the transcription machinery.

## Discussion

Our results identify the human FACT complex as a key facilitator of cohesin translocation on chromatin and, consequently, of cohesin-dependent genome folding. We show that FACT and cohesin co-occupy a substantial fraction of genomic sites (Fig.1 a-e) and physically interact (Fig.1 f), consistent with a conserved functional relationship from yeast to mammals. Acute depletion of the FACT subunit SUPT16H does not reduce the overall levels of chromatin-bound cohesin (Fig.2 c) but profoundly alters its distribution: cohesin is depleted from loop anchors and TAD boundaries (Fig.2 e; Fig.3 e-g) while accumulating across transcribed gene bodies (Fig. 4 a-f). These changes are accompanied by fewer and weaker loops, reduced boundary insulation, and diminished stripe intensity (Fig.3 c-h). Functionally, cohesin accumulation within gene bodies correlates with transcriptional decline (Fig.5 a), and co-depletion of RAD21 restores transcription at affected loci (Fig.5 b,c). Together, these findings support a model in which FACT promotes productive cohesin translocation, enabling efficient loop extrusion while preventing aberrant cohesin retention within actively transcribed regions.

A central implication of our data is that cohesin movement through chromatin *in vivo* is strongly constrained by chromatin context and therefore requires factors that facilitate its progression. While *in vitro* studies have shown that SMC complexes can bypass large obstacles under simplified conditions (Pradhan et al., 2022), chromatin *in vivo* presents a dense and heterogeneous landscape of nucleosomes and DNA-bound proteins which might present a challenge (Davidson et al., 2016; Stigler et al., 2016). Indeed, RNA polymerase II (RNAPII) acts as a moving barrier that redistributes cohesin (Banigan et al., 2022; Jeppsson et al., 2022), and MCM complexes function as chromatin-bound obstacles that restrict cohesin translocation (Dequeker et al., 2022). Our data place FACT within this framework as a permissive factor that alleviates these constraints. Through its established role as a histone chaperone, FACT likely facilitates cohesin movement by transiently destabilizing nucleosomes, thereby lowering the effective barrier to translocation.

Consistent with this view, SUPT16H depletion weakens loop signals and TAD insulation without substantially altering the genomic distance scaling of contacts (Fig.3 d,f). This behaviour mirrors perturbations that reduce the efficiency of cohesin reaching anchoring sites rather than abolishing extrusion altogether (Dequeker et al., 2022; Wutz et al., 2017). We therefore propose that, in the absence of FACT, cohesin remains capable of loading and limited movement, but fails to efficiently traverse chromatin and reach convergent CTCF sites. As a result, focal contacts weaken and domain insulation is reduced, even though the intrinsic scale of loop extrusion is largely preserved.

The redistribution of cohesin into gene bodies upon FACT depletion provides further mechanistic insight. Under normal conditions, cohesin positioning reflects a dynamic balance between extrusion, release, stalling at CTCF, and interactions with the transcription machinery (Banigan et al., 2022; Busslinger et al., 2017; Jeppsson et al., 2022; Lengronne et al., 2004). Elongating RNAPII has been proposed to act as a moving barrier that can reposition or displace cohesin along gene bodies (Banigan et al., 2022; Jeppsson et al., 2022). Our data extend this model by showing that this process critically depends on FACT. In the presence of FACT, efficient RNAPII elongation and nucleosome remodelling facilitate both transcription and cohesin translocation. In contrast, FACT depletion impairs polymerase processivity and increases the chromatin barrier, leading to cohesin retention within gene bodies, particularly at highly transcribed loci.

Importantly, this redistribution has functional consequences for transcription. Genes that accumulate cohesin upon SUPT16H depletion exhibit a pronounced reduction in nascent transcription, and this defect is rescued by simultaneous depletion of RAD21. These results demonstrate that transcriptional impairment is not solely a direct consequence of FACT loss but is substantially mediated by aberrant cohesin accumulation. Thus, while RNAPII can act as a barrier that shapes cohesin distribution, excessive cohesin retention can in turn impede transcription (Fig.5 c), revealing a reciprocal interplay between these processes.

Our Hi-C analysis further indicates that cohesin retained within gene bodies is not productively engaged in loop extrusion (Fig.3). Rather than enhancing local interactions, cohesin accumulation is associated with reduced intra-genic contacts and loss of genebody insulation. This suggests that cohesin becomes trapped in a non-productive state in which it remains chromatin-bound but is impaired in its ability to extrude DNA and generate higher-order structure. This interpretation is consistent with the global weakening of loops and stripes, pointing to a defect in cohesin dynamics rather than its abundance.

Our findings also inform the interpretation of NIPBL localisation. Although NIPBL is enriched at promoters and often interpreted as marking cohesin loading sites, its redistribution across gene bodies upon FACT depletion argues that it can remain associated with cohesin after loading. This behaviour is more consistent with a role within the active or stalled extrusion machinery than with a purely loading-specific function.

More broadly, our study highlights histone chaperones as key regulators of genome architecture. Whereas CTCF provides sequence-specific barriers and transcription complexes impose dynamic constraints, FACT appears to promote cohesin mobility by modulating chromatin structure. These observations place chromatin remodelling at the core of loop extrusion *in vivo*, suggesting that the efficiency of cohesin translocation is determined not only by cohesin-associated factors such as NIPBL and WAPL, but also by the physical properties of the chromatin fibre itself.

Several questions remain. The molecular basis of the FACT-cohesin interaction and how nucleosome reorganization is coupled to cohesin translocation remain to be defined. It will also be important to determine whether cohesin complexes that accumulate in gene bodies are physically stalled, compositionally altered, or topologically constrained. Finally, although our data support a role for cohesin accumulation in transcriptional repression, the precise mechanism; whether steric interference with RNAPII, disruption of chromatin reassembly, or altered DNA topology; remains to be established.

In summary, we propose that FACT promotes cohesin translocation through chromatin, thereby sustaining loop extrusion, loop anchoring, and domain insulation. In its absence, cohesin becomes trapped within gene bodies, uncoupled from productive genome folding, and inhibitory to transcription. FACT thus emerges as a central coordinator of chromatin dynamics, linking nucleosome remodelling to cohesin function and the integration of 3D genome organization with transcriptional output.

## Methods

### Cell lines

Human HCT116 cell lines were used in this study. Cells expressing CMV-OsTIR1 and CMV-OsTIR1(F74G) integrated at the AAVS1 locus were described previously (Natsume et al., 2016; Yesbolatova et al., 2020). Cells were maintained in McCoy’s 5A medium (modified) supplemented with 10% FBS, 1× antibiotic-antimycotic and 2 mM L-glutamine. To generate SUPT16H-AID cells, HCT116 CMV-OsTIR1 cells were co-transfected with a CRISPR plasmidb(pX330-SUPT16H-C) targeting 5’-AG-GAAGTAACTTCTGAACTT-3’ and a donor plasmid (SUPT16H-mAID-mClover) harbouring 500-bp homology arms and a hygromycin-resistant marker following the published protocol (Yesbolatova et al., 2019). Following selection with 100 μg/mL HygroGold, single-cell clones were isolated. Homology-directed integration of the mAC tag at both SUPT16H alleles was confirmed by genomic PCR and Western blotting. To generate RAD21-mAID-mCherry, SUPT16H-mAID-mClover cell line, we initially fused mAID-mClover to the C-terminus of SUPT16H in the HCT116 cells expressing CMV-OsTIR1(F74G) using the same CRISPR and donor plasmids described above. Subsequently, we co-transfected with a donor plasmid (pX330-RAD21-C) targeting 5’ CCAAGGTTCCATATTATATA3’ and a donor plasmid (RAD21-mAID-mCherry) harbouring 500-bp homology arms and a neomycin-resistant marker. Following transfection, cells were selected with 100 μg/mL G418. Single-cell clones were isolated and correct integration of RAD21-mAID-mCherry was confirmed by genomic PCR and Western blotting.

HCT116 cells constitutively expressing CMV-OsTIR1 were pre-treated with 200 μM auxinole, an OsTIR1 inhibitor that suppresses basal degradation of mAID-fused protein, prior to auxin addition. Degradation of mAID-fused protein was then induced by replacing the medium with fresh medium containing 500 μM indole-3-acetic acid (Auxin, IAA) for 6 hours. Auxinole and IAA were dissolved in DMSO to 200 mM and 500 mM stock solutions, respectively, and stored at-20°C

In HCT116 cells constitutively expressing OsTIR1(F74G) degradation of mAID-fused proteins was induced by the addition of 1 µM 5-phenyl-indole-3-acetic acid (5-Ph-IAA) dissolved in DMSO (1 mM stock, stored at-20°C) directly to the culture medium for 6 hours.

### Microscopy

HCT116 cells expressing SUPT16H-mAID-mClover were used to monitor auxin-induced SUPT16H degradation. Cells were treated with auxin or auxinole as control and imaged over time.

For live-cell analysis, cells were seeded in 48-well plates and imaged using an IncuCyte Zoom system (Essen Bioscience). Phase contrast and green fluorescence images were acquired at regular intervals throughout the experiment. Quantification of RAD21-Clover levels was performed using the built-in IncuCyte analysis software by measuring the Green Integrated Intensity normalized to Phase Confluence, which accounts for differences in cell density. Fluorescence values were normalized to the initial time point. For higher-resolution imaging, cells were seeded in µ-Slide 8 Well chambers (ibidi) and imaged under controlled environmental conditions. Imaging was performed using an Olympus IX70 epifluorescence microscope equipped with a Lumencor Spectra LED light source, a Hamamatsu Orca Flash 4.0 V2 camera, and a 40× objective (NA 0.95, UPlanSApo). A stage-top incubation chamber was used to maintain 37C and 5% CO₂ throughout the experiment.

### Western blot

Protein extraction was done by lysing the cells with 20% TCA. Proteins were precipitated with TCA 7.5% and centrifuging at 15000 rpm for 10 minutes at 4 °C. Then the pellet was resuspended in Laemmli buffer 1.5X. Western blots were resolved in 7.5% SDS-PAGE gels. Proteins were transferred to polyvinylidene fluoride (PVDF) membranes using the TE70X semidry blotter system (Hoefer). The antibodies used were anti-SMC1 (AbCam, ab9262), anti-SUPT16H (Santa Cruz, sc-377028), anti-CTCF (AbCam, ab70303) anti-tubulin (Santa Cruz, sc-5286) and anti-H3 (AbCam, ab18521). Blots were incubated with the ECL Prime Western blotting detection reagent (GE Healthcare). Blots were developed by exposure in an ImageQuant LAS 4000 mini machine (GE Healthcare).

### Chromatin binding assay

7 × 10⁶ HCT116 cells were harvested, washed in ice-cold PBS, and resuspended in extraction buffer (50 mM HEPES-KOH pH 7.5, 100 mM KCl, 2.5 mM MgCl₂, 10 mM DTT, protease inhibitors) supplemented with 0.25% Triton X-100. Cells were incubated on ice for 5 min to allow lysis.

An aliquot was retained as whole-cell extract (WCE), while the remaining lysate was layered over a sucrose-containing buffer cushion (50 mM HEPES-KOH pH 7.5, 100 mM KCI, 2.5 mM MgC)z, 10 mM OTT, 0.25 %Tx-100, 30% Sucurose) and centrifuged at 12,000 rpm for 10 min at 4°C to separate soluble (SN) and chromatin-bound (CP) fractions. The supernatant was collected as the soluble fraction, and the chromatin pellet was washed once with extraction buffer and resuspended.

All fractions were mixed with SDS sample buffer, boiled, and analyzed by SDS-PAGE and immunoblotting.

### Protein co-immunoprecipitation

7*10^6^ HCT116 cells were trypsinized and washed in cold water and resuspended in 200 μl of ice-cold buffer A (50 mM HEPES, 150 mM KCl, 1.5 mM MgCl2, 0.5 mM DTT and 0.5% Triton X-100 (pH 7.5) supplemented with complete protease inhibitor cocktail tablets (Roche)). Cells were lysed passing cells trough a 25 gauge needle 10 times on ice. Extracts were maintained on ice for 2 min after each cycle. Cell extracts were centrifuged for 10 minutes at 12000 r.p.m at 4C and the supernatant incubated with protein G Dynabeads (Invitrogen) bound to anti-SMC1 antibody (AbCam, ab9262) or anti-SUPT16H antibody (AbCam, ab204343) for 2 hours at 4 °C. Finally, beads were washed five times in washing buffer (10 mM Tris-Cl pH 7.5, 150 mM NaCl, 0.5% Triton) and unbound from the antibody by incubating at 37°C for 4 min in SR buffer (2% SDS, 0.125 M Tris-Cl pH 6.8). Immunoprecipitated proteins were mixed with SS buffer (5% saccharose, 0.0125% bromophenol blue) and run in an SDS-PAGE gel.

### Chromatin immunoprecipitation sequencing (ChIP-seq)

ChIP-seq was performed essentially as described previously (Canela et al., 2017) with some modifications. Briefly, 20 million HCT116 cells were fixed with 1% formaldehyde for 10 min at 37°C and fixation was quenched with 125 mM glycine. For mouse spike-in experiments, 1.5 million of mouse Abelson pre-B cells (7.5%) arrested in G1 with imatinib, were added prior to fixation. Chromatin was prepared in RIPA buffer, sonicated using a Branson sonifier, and immunoprecipitated overnight using Dyna-beads Protein A coupled to the indicated antibodies. For experiments using Drosophila spike-in normalization, Drosophila chromatin (Active Motif, Cat. 53083) and spike-in antibody (Active Motif, Cat. 61686) were added before immunoprecipitation. Following sequential high-salt washes, crosslinks were reversed, DNA was purified using phenol:chloroform:isoamilic following ethanol precipitation, and sequencing libraries were prepared using standard Illumina protocols. Libraries were sequenced as single-end 75-bp reads on an Illumina NextSeq 500 platform. Antibodies used were anti-RAD21 (Abcam, ab992), anti-CTCF (Millipore, 07-729), anti-NIPBL (Bethyl, A301-779) and anti-GFP (Abcam, ab290).

### Nascent RNA-seq

Nascent RNA was labelled by incubating cells with 0.5 mM 5-ethynyl uridine (EU) for 30 min, after which cells were harvested, washed in cold PBS and snap-frozen. Total RNA was extracted using TRIzol/chloroform followed by isopropanol precipitation. For each sample, 1 μg of total RNA was depleted of ribosomal RNA using the NEB-Next rRNA Depletion Kit (Human/Mouse/Rat) and purified with RNAClean XP beads. EU-labelled nascent RNA was then biotinylated by click chemistry using the Click-iT Nascent RNA Capture Kit (Thermo Fisher) and captured on Dynabeads MyOne Streptavidin T1 beads. Bead-bound RNA was reverse transcribed with SuperScript VILO, followed by second-strand synthesis in the presence of dUTP to preserve strand information. Double-stranded cDNA was end-repaired, A-tailed, ligated to TruSeq adapters, treated with uracil-DNA glycosylase, and PCR-amplified with KAPA HiFi polymerase. Libraries were size-selected from 200-400 bp, quantified by qPCR, and sequenced as single-end 75-bp reads on an Illumina platform.

### ChIP-seq and Nascent RNA-seq analysis

Nascent RNA-seq reads were mapped to the human (GRCh37/hg19) genome using Bowtie2 (v2.5.1-1) (Langmead and Salzberg, 2012) with the default parameters. ChIP-seq reads were aligned to combined human (GRCh37/hg19) and Drosophila (dm6), or human (GRCh37/hg19) and mouse (GRCm38/mm10), reference genomes using Bowtie2 (v2.5.1-1)(Langmead and Salzberg, 2012) with default parameters. The resulting human-aligned BAM files were converted to bedGraph files using bedtools (Quinlan and Hall, 2010) scaling to reads per million mapped human reads and multiplying by the corresponding spike-in calibration factor. BedGraph files were then converted to bigWig format using bedGraphToBigWig (Karolchik et al., 2004). Genomic profiles were visualized using the UCSC Genome Browser. For spike-in calibrated samples, calibration factors were calculated multiplying the ratio of spike-in reads/human reads in the input and the ratio of human reads/ spike-in reads in the sample. For the SUPT16H-GFP biological replicate 1 samples, for which no matched spike-in input was available, calibration was performed using the relative spike-in abundance between ChIP libraries after normalization to total mapped reads.

Peaks were identified using MACS (v1.4.3) (Zhang et al., 2008) with default parameters and peaks later were filtered for >3-fold enrichment of RPKM over input. RPKM values were multiplied by the corresponding spike-in calibration factor. Downstream analyses were performed using bedtools (v2.31.1) (Quinlan and Hall, 2010) and R (v4.3.2).

The refseq gene annotations (hg19/GRCh37), taken from the UCSC database (Karolchik et al., 2004) were used to obtain the Transcription start sites (TSS), transcription end sites (TES) coordinates. All transcripts’ isoforms for a given gene were merged into a single gene coordinate with the outer TSS and TES. Genes at the chrM, chrUn, chr random and haplotype chromosomes were excluded from the analysis, and only genes with length longer than 2 kb were retained. For genebody quantification, ChIP-seq read densities were calculated as reads per kilobase per million mapped reads (RPKM) across each gene body after excluding RAD21 peak intervals identified in control (auxinole) samples. ChIP-seq RPKM values were multiplied by the corresponding spike-in calibration factor. For nascent RNA-seq, RPKM values were calculated across the entire gene body. These gene-body RPKM values were used for all gene scatter plot analyses. For nascent RNA-seq quantification, reads mapping to the same 5’ genomic coordinate were collapsed into a single count before calculating read densities to reduce the contribution of PCR bias. ChIP-seq analyses were performed using all mapped reads.

Heatmaps were generated using the R package pheatmap (1.0.13). Common RAD21 peaks in control (auxinole) samples were centered on the peak summit, and read start densities per million mapped reads were quantified within a +/-1 kb interval around each summit. Peak intervals were sorted in decreasing order according to the RAD21 signal intensity in the control condition, and the top 20,000 peaks were plotted. The same intervals and ordering were subsequently used to quantify signal and generate heatmaps for the additional ChIP-seq control datasets. Input color scales were matched to those used for NIPBL and SUPT16H.

For aggregate profiles spanning genomic intervals, the distance between two positions was divided into 200 equal-sized bins. Depending on the analysis, these positions corresponded to TSS and TES, pairs of CTCF-bound RAD21 peaks (CTCF+ RAD21+), pairs of Hi-C loop anchors, or pairs of Hi-C TAD boundaries. Upstream and downstream flanking regions were represented by 50 bins. For nascent RNA-seq analyses, flanking regions corresponded to fixed 2 kb windows were used to preserve genomic distance information at promoters and transcription termination regions. For ChIP-seq analyses, the bin size of flanking regions was the same as genomic interval bins, allowing comparison of signal distributions across genomic features of variable size.

Reads overlapping each bin were quantified using bedtools intersect and bedtools groupby. Read densities were calculated for each bin and normalized by bin length and total mapped reads to generate reads per kilobase per million mapped reads (RPKM) values multiplied by the spike-in calibration factor. Aggregate profiles were computed by averaging signal across all genes or intervals in the indicated set and plotted using smooth.spline function in R.

For RAD21 and NIPBL metagene analysis comparing SUPT16H depletion, the intervals corresponding to RAD21 and NIPBL peaks respectively within the gene body in the control (auxinole) samples were excluded from quantification. Additionally, gene body intervals were classified into four quartiles according to the average RPKM of the control (auxinole) nascent RNA-seq libraries.

For Hi-C TAD boundaries and loop anchors identified by HOMER common in the control (auxinole) samples were expanded on each side 20 kb for TAD boundaries and 10 kb for for loops, and genomic intervals were defined by the outermost coordinates. Additionally, for the analysis shown in Supplementary Fig. 4C, TADs were classified into four quartiles according to the transcriptional activity of the genes contained within each expanded TAD.

For RAD21 ChIP-seq aggregate profiles associated with putative cohesin loading sites (RAD21+ NIPBL+ CTCF-), RAD21 signal was quantified from each loading-site peak to the nearest RAD21+ NIPBL-CTCF+ peak located within 200 kb upstream or downstream. Loading-site intervals and RAD21+ NIPBL-CTCF+ peak intervals were excluded from the quantified region. Signal was quantified at single base-pair resolution and subsequently binned into 1 kb intervals. Upstream and downstream intervals were reoriented and collapsed into a common direction with the loading site positioned at the origin of the plot. RPKM values in each bin were multiplied by the corresponding spike-in calibration factor, and aggregate profiles were computed by averaging the signal across all intervals. Curves were plotted in R using the smooth.spline function.

### Hi-C libraries

Hi-C was performed as previously described for two conditions: HCT116 and HCT116 cells with depleted SUPT16H, with two biological replicates per condition. Per replicate, 5 million cells were lysed in a lysis buffer (30 minutes on ice) and digested with DpnII (NEB) overnight at 37°C while rotating (950 rpm). Restriction overhangs were filled in with Klenow (NEB) using biotin-14-dATP (Jena Bioscience), and ligation was performed in the ligation buffer for 4 hours at 16°C (T4 DNA ligase; Life Technologies). After overnight de-crosslinking at 65°C, the ligated DNA was tagmented to produce fragments of 300-700 bp. Ligation products were isolated using MyOne C1 streptavidin beads (Life Technologies), followed by washing with Wash&Binding buffer and nuclease-free water. Isolated Hi-C ligation products on the beads were then used directly for PCR amplification (5 PCR cycles), and the final Hi-C library was purified with AMPure XP beads (Beckman Coulter). The final libraries were paired-end sequenced (50 bp) on an Illumina HiSeq 2500 machine at a sequencing depth of 600 million paired reads per replicate.

### Generation and normalization of contact maps

Hi-C data were processed using HiC-Pro (v3.1.0) with default parameters unless otherwise specified (Servant et al., 2015). Reads were aligned to the hg19 reference genome using Bowtie2, and alignments with mapping quality <10 were discarded. PCR duplicates, multi-mapped reads, and singleton pairs were removed. Reads were assigned to restriction fragments generated using the DpnII restriction enzyme. Contact matrices were generated at multiple resolutions (1, 5, 10, 25, 100 and 500 kb) and stored in upper-triangular format. Normalization was performed using the ICE algorithm with a maximum of 100 iterations and filtering of low-coverage bins. HiC-Pro output matrices were subsequently converted to.cool format using HiC-Pro utilities and further normalized and processed using the cooler toolkit.

### Matrices visualization

Hi-C contact matrices were visualized using HiGlass (Kerpedjiev et al., 2018). Multiresolution.cool files were generated from HiC-Pro output using the hicpro2higlass.sh utility and loaded into a locally run HiGlass instance running in Docker. Data were visualized through the HiGlass web interface with continuous zooming and synchronized multi-scale views, enabling inspection of chromatin interactions across genomic regions and resolutions.

### Compartment detection and compartment strength on Hi-C contact maps

Compartment detection was performed on Hi-C contact maps at 100-kb resolution using cooltools (Open2C et al., 2024). Balanced contact matrices in.cool format were used to compute cis eigenvectors, and the first eigenvector (E1) was used to assign A/B compartments across each chromosome. For compartment strength analysis, expected contact frequencies were computed and genomic bins were stratified according to E1 values. Saddle matrices were then calculated to quantify preferential interactions between compartments and visualized as observed/expected contact enrichments.

### Contact probability as a function of the genomic distance P(s)

Contact probability as a function of genomic distance (P(s)) was calculated from 10-kb balanced Hi-C matrices in.cool format using cooltools (Open2C et al., 2024). Analyses were performed separately for each replicate. Smoothed and aggregated P(s) curves were generated and comparison between conditions, replicate curves were grouped as control (+ auxinole) or SUPT16H depleted (+auxin), and averaged at each genomic distance. Distances below 12 kb were excluded from downstream analysis and plotting. The slope of the P(s) curve was calculated as the derivative in log-log space using np.gradient on the averaged smoothed contact frequencies.

### Detection of TADs and Loops

Topologically associating domains (TADs) and chromatin loops were identified using HOMER (Heinz et al., 2018). HiC-Pro validPairs files were converted into HOMER-compatible Hi-C summary. These reformatted files were then used to generate HOMER tag directories with makeTagDirectory using the-format HiCsummary option. The resulting tag directories were used as input for findTADsAndLoops.pl to detect TADs and loops at at 3-kb resolution with a 15-kb window size (-res 3000-window 15000) using the hg19 genome.

### Detection of stripes

Architectural stripes were identified from Hi-C contact maps using Stripenn (v1.1.65) (Yoon et al., 2022). Balanced contact matrices in.cool format were analyzed at 5 kb resolution using the compute function with KR normalization. Stripe detection was performed genome-wide using default parameters. The resulting stripe coordinates, along with associated metrics including length, width, mean signal intensity, p-value, and stripiness score, were used for downstream analyses and comparisons between conditions.

### Aggregate plots

Aggregate Hi-C analyses (pile-ups) of loops, TADs, stripes and gene bodies were performed using coolpup.py on balanced contact matrices in.cool format. Genomic coordinates corresponding to loops, TAD boundaries, or stripe-associated regions were used as input, and interaction submatrices were extracted and averaged to generate aggregate interaction profiles. Pile-ups were normalized to distance-dependent expected contact frequencies to account for the decay of contact probability with genomic distance. Resulting aggregate matrices were visualized to quantify average enrichment of loop interactions, domain boundaries, stripe-like features and gene bodies interactions across conditions.

## Data availability

Full details of ChIP-seq libraries and raw sequencing data are available at GEO under accession number GSE335764, nascent RNA-seq data under accession number GSE335766, and Hi-C data under SRA accession PRJNA1474361. All other data that support the conclusions are available from the corresponding authors on reasonable request.

