## Supplementary figures for "Human FACT complex coordinates cohesin and transcription on chromatin"

### Figure S1

a

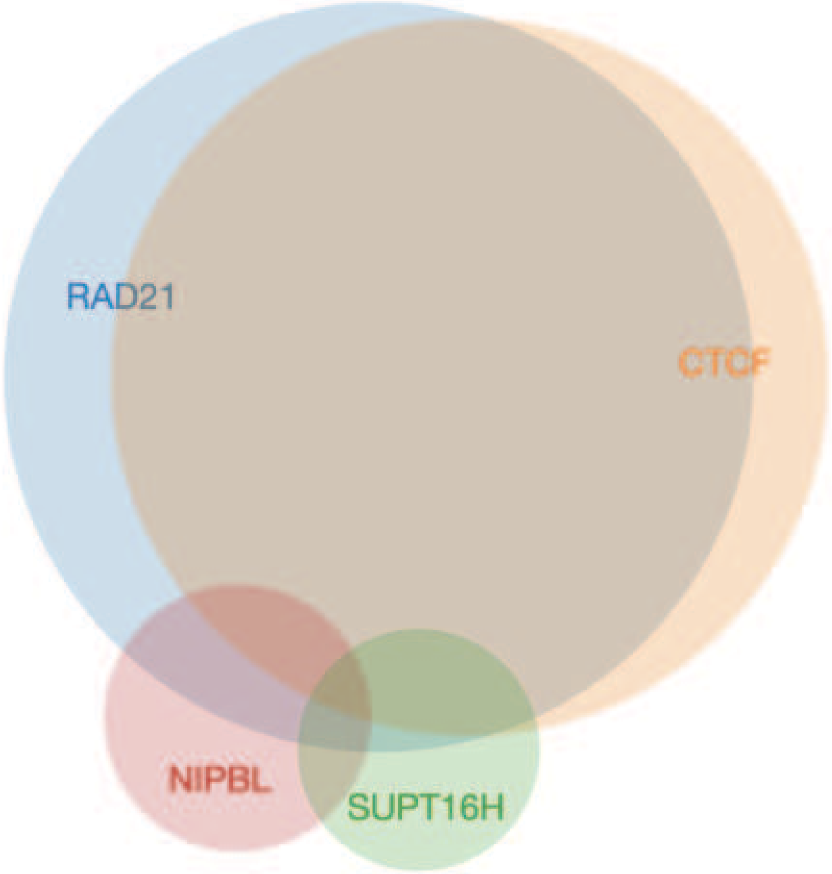


| | Nº of peaks |
|---|---|
| RAD21 | 44765 |
| CTCF | 40971 |
| SUPT16H | 4666 |
| NIPBL | 5680 |
| RAD21^CTCF | 36018 |
| RAD21^SUPT16H | 2258 |
| RAD21^NIPBL | 2722 |
| CTCF^SUPT16H | 1760 |
| CTCF^NIPBL | 1344 |
| SUPT16H^NIPBL | 821 |

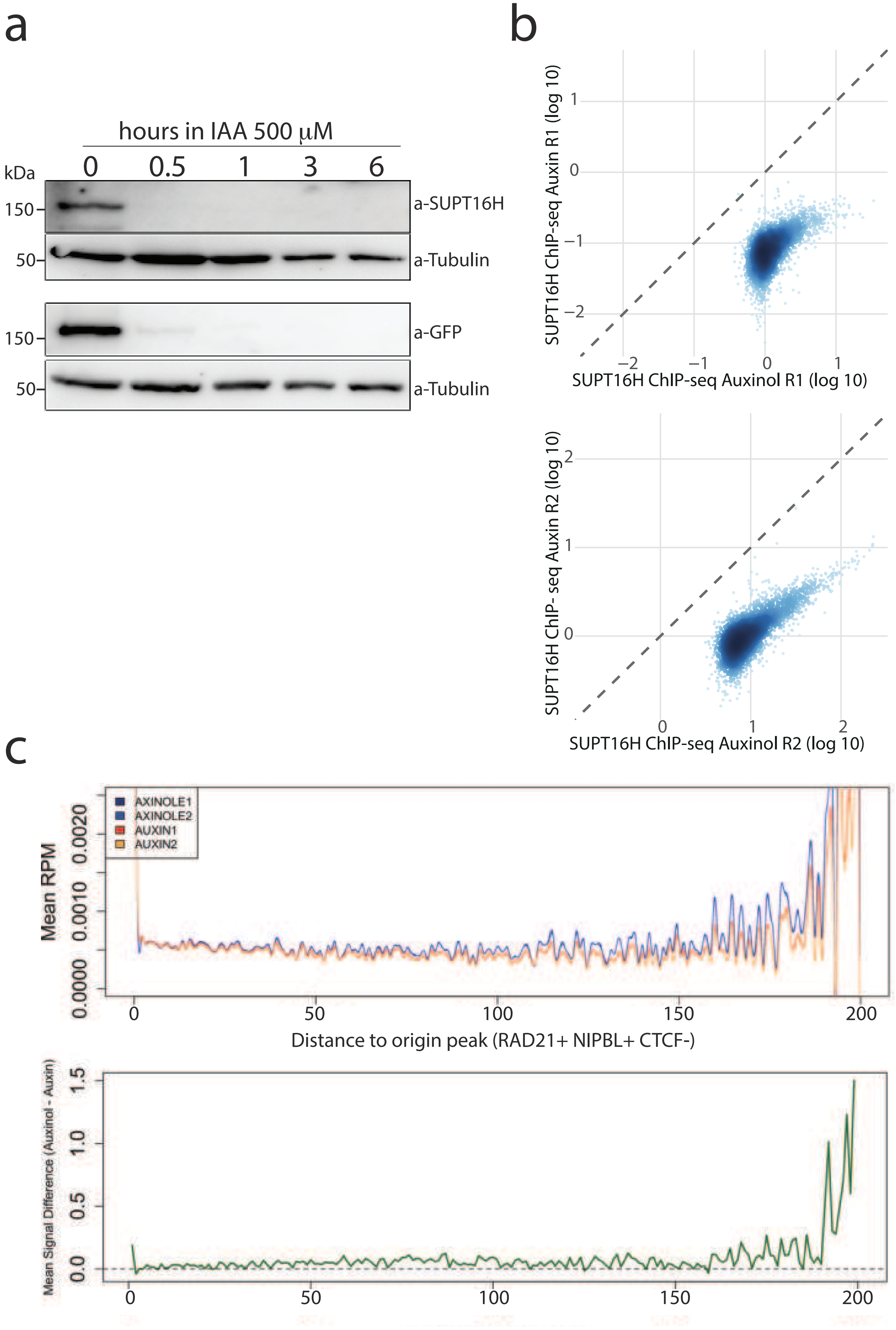
a
hours in IAA 500 μM
0
0.5
1
3
6
kDa
150
50
150
50
a-SUPT16H
a-Tubulin
a-GFP
a-Tubulin
b
SUPT16H ChIP-seq Auxin R1 (log 10)
SUPT16H ChIP-seq Auxinol R1 (log 10)
SUPT16H ChIP- seq Auxin R2 (log 10)
SUPT16H ChIP-seq Auxinol R2 (log 10)
c
AXINOLE1
AXINOLE2
AUXIN1
AUXIN2
Mean RPM
Distance to origin peak (RAD21+ NIPBL+ CTCF-)
Mean Signal Difference (Auxinol - Auxin)

### Figure S3

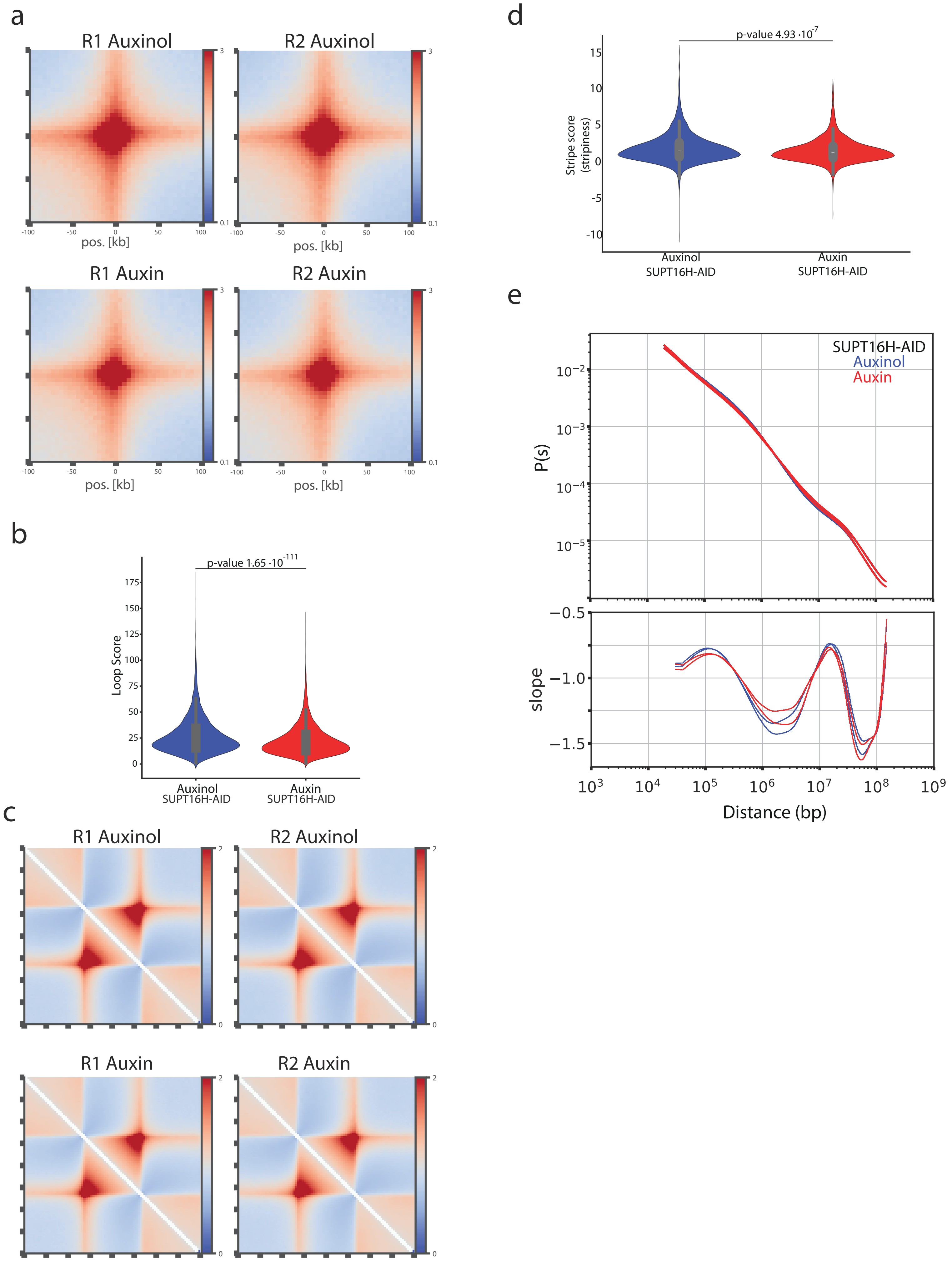
a
R1 Auxinol
R2 Auxinol
R1 Auxin
R2 Auxin
pos. [kb]
-100
-50
0
50
100
3
0.1
b
p-value 1.65 ·10-111
Loop Score
175
150
125
100
75
50
25
0
Auxinol
SUPT16H-AID
Auxin
SUPT16H-AID
c
R1 Auxinol
R2 Auxinol
R1 Auxin
R2 Auxin
2
0
d
p-value 4.93 ·10-7
Stripe score
(stripiness)
15
10
5
0
-5
-10
Auxinol
SUPT16H-AID
Auxin
SUPT16H-AID
e
SUPT16H-AID
Auxinol
Auxin
P(s)
10−2
10−3
10−4
10−5
slope
−0.5
−1.0
−1.5
10³
10⁴
10⁵
10⁶
10⁷
10⁸
10⁹
Distance (bp)

### Figure S4

a

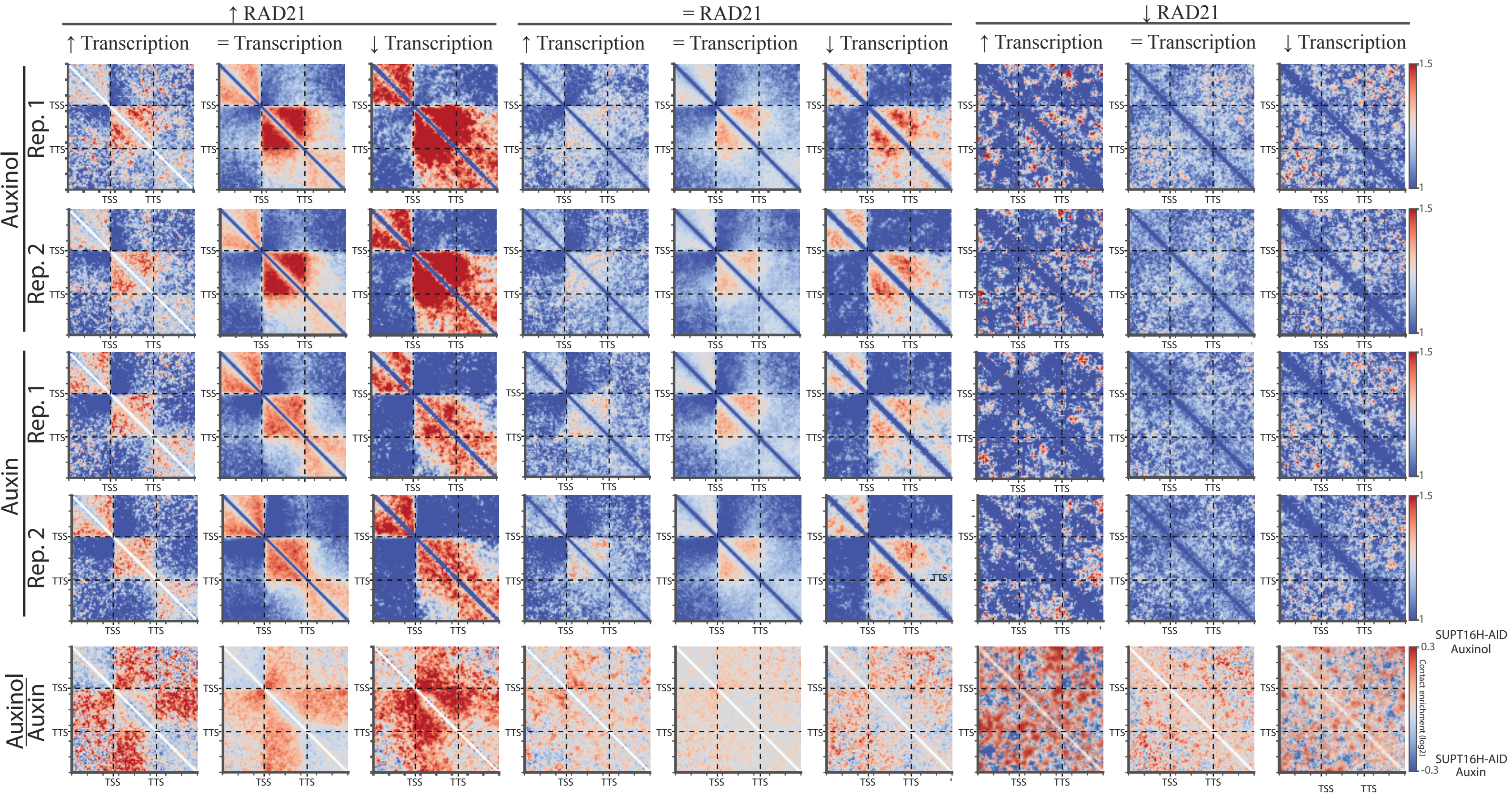

↑ RAD21
= RAD21
↓ RAD21
↑ Transcription
= Transcription
↓ Transcription
Auxinol
Auxin
Rep. 1
Rep. 2
Auxinol / Auxin
TSS
TTS
1.5
1
0.3
-0.3
Contact enrichment (log2)
SUPT16H-AID Auxinol
SUPT16H-AID Auxin


b

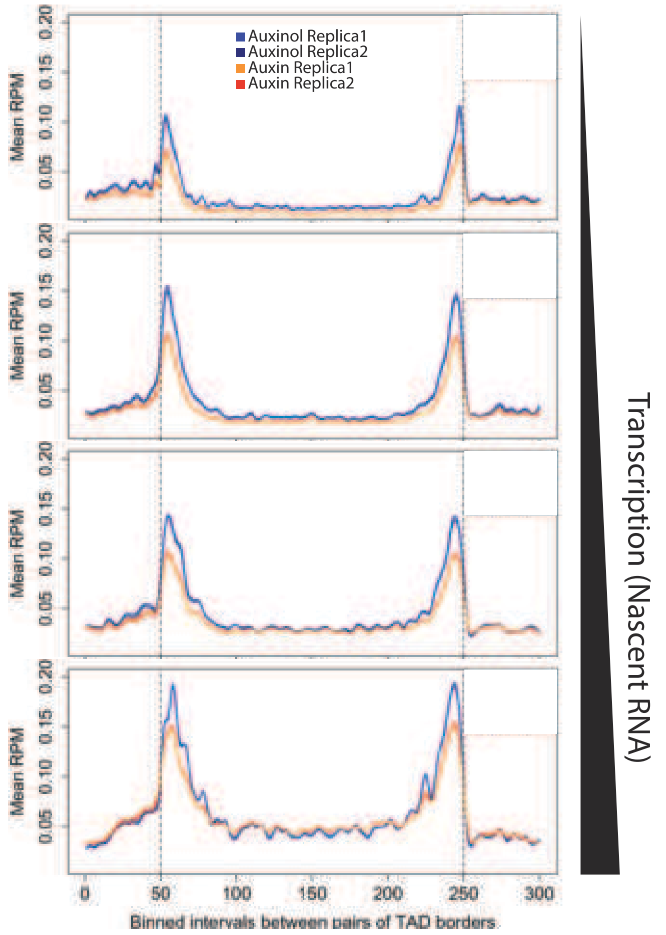

Auxinol Replica1
Auxinol Replica2
Auxin Replica1
Auxin Replica2
Mean RPM
0.05
0.10
0.15
0.20
0
50
100
150
200
250
300
Binned intervals between pairs of TAD borders
Transcription (Nascent RNA)

### Figure S5

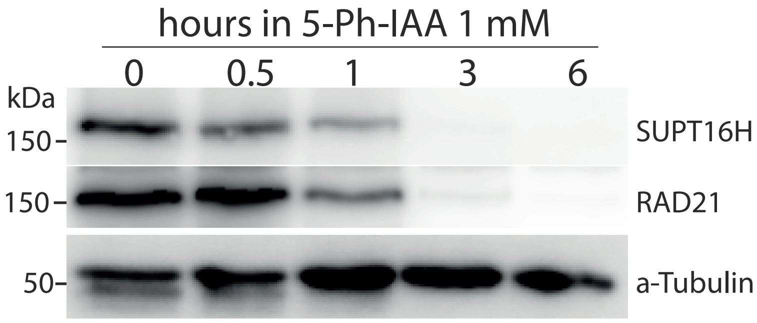
hours in 5-Ph-IAA 1 mM
0
0.5
1
3
6
kDa
150
150
50
SUPT16H
RAD21
a-Tubulin

### Supplementary Information

Human FACT complex coordinates cohesin and transcription on chromatin

Jonay Garcia-Luis, Andres Canela, Valeriya Malysheva, Toyoaki Natsume, Mohammad M Karimi, Masato T. Kanemaki, Mikhail Spivakov & Luis Aragon

**Figure S1 | Genome-wide co-localization of FACT, cohesin, and associated factors. a,** Venn diagram showing the overlap between ChIP-seq peaks for RAD21, CTCF, SUPT16H, and NIPBL in HCT116 cells. A substantial fraction of SUPT16H peaks co-localizes with RAD21, indicating widespread co-occupancy of FACT and cohesin across the genome. Overlaps between all factor combinations are quantified in the table on the right.

**Figure S2 | Rapid depletion of SUPT16H alters cohesin distribution without affecting bulk chromatin association. a,** Western blot showing time-course degradation of SUPT16H-AID-mClover following auxin (IAA, 500 μM) addition in HCT116 cells. SUPT16H is detected with a-SUPT16H specific antibody or a-GFP antibody. Signals decrease after 30 min. of auxin treatment **b,** Scatter plot comparing SUPT16H ChIP-seq signal in SUPT16H depleted (+auxin) versus control (+auxinole) conditions, illustrating global depletion of SUPT16H upon auxin treatment. **c,** Scatter plots comparing RAD21 ChIP-seq signal at all RAD21 peaks (n = 59,409) between control (+auxinole) and SUPT16H-depleted (+auxin) conditions for biological replicates R1 and R2. **d,** Metaprofile of calibrated RAD21 ChIP-seq signal between putative cohesin loading sites ($RAD21^{+}$ $NIPBL^{+}$ $CTCF^{-}$) and the closest RAD21+ NIBL- CTCF+ peak located withing 200kb upstream or downstream. Loading and RAD21+ NIBL- CTCF+ peak intervals were excluded from the quantified regions. Signal was quantified at single-base resolution, binned into 1 kb intervals and plotted after collapsing upstream and downstream regions into the same orientation. Top: mean RPKM signal for biological replicates in control (+auxinole) and SUPT16H depleted (+auxin) conditions. Bottom: mean signal difference (Auxinole - Auxin). SUPT16H depletion leads to reduced RAD21 enrichment distal to cohesin loading sites, consistent with impaired cohesin translocation along chromatin.

**Figure S3 | SUPT16H depletion weakens cohesin-dependent chromatin architecture without altering global scaling. a,** Aggregate peak analysis (APA) of loop signals from Hi-C data in control (+auxinole) and SUPT16H depleted (+auxin) cells across two biological replicates (R1, R2). Loop-associated contact enrichment is reduced upon SUPT16H depletion. **b,** HOMER-based quantification of loop strength shows a significant decrease upon auxin treatment, consistent with weakened cohesin-mediated looping. Violin plots show the distribution of loop scores for Control (+Auxinole) and SUPT16H-depleted (+Auxin) conditions, calculated as the mean of two replicates. The width represents data density, the white line indicates the median, and the grey box shows the interquartile range (Q1-Q3). p-value calculated using the Mann-Whitney U test (two-sided). **c,** Aggregate TAD boundary analysis (pile-ups) from Hi-C contact maps across two biological replicates in control and SUPT16H-depleted conditions (R1, R2). Boundary-associated contact patterns are weakened upon FACT depletion. **d,** Quantification of stripe strength using Stripenn reveals a significant reduction in stripe scores following SUPT16H degradation, indicating impaired cohesin processivity. Violin plots show the distribution of stripiness scores for Control (+Auxinole) and SUPT16H-depleted (+Auxin) conditions. The width represents data density, the white line indicates the median, and the grey box shows the interquartile range (Q1-Q3). p-value calculated using the Mann-Whitney U test (two-sided) **e,** Contact probability as a function of genomic distance, P(s), for control and SUPT16H-depleted cells (top), and corresponding slope analysis (bottom) across two biological replicates.

**Figure S4 | Transcription-dependent cohesin and NIPBL accumulation across gene bodies and altered chromatin organisation upon FACT depletion. a,** Metagene analysis of calibrated RAD21 ChIP-seq signal across gene bodies. Genes were divided into four quartiles (Q1-Q4) based on nascent RNA levels, from lowest (Q1) to highest (Q4) transcriptional activity. The distance between the transcription start site (TSS) and transcription end site (TES) was divided into 200 equal bins, with 50 flanking bins on each side; genes shorter than 2,000 bp were excluded. Data are shown for two biological replicates under control (+Auxinole, blue) and SUPT16H-depleted (+Auxin, red) conditions. Upon SUPT16H depletion, RAD21 accumulates broadly across gene bodies in a transcription-dependent manner, with the greatest increase observed in the most highly transcribed genes (Q4). **b,** As in (a), for calibrated NIPBL ChIP-seq signal (one replicate shown). NIPBL displays a similar but more restricted increase in gene-body enrichment upon SUPT16H depletion. **c,** Hi-C pile-up matrices centred on gene bodies (TSS to TES) stratified by changes in RAD21 enrichment (increased, unchanged, decreased) and transcriptional response (increased, unchanged, decreased) upon SUPT16H depletion. Data are shown for two biological replicates under control (+Auxinole) and SUPT16H-depleted (+Auxin) conditions, as well as the Auxinole/Auxin dratio of both replicates combined. Cohesin accumulation is associated with reduced intra-genic contact and weakened gene-body insulation independently of transcriptional changes. **d,** Metaplot of calibrated RAD21 ChIP-seq signal across TADs stratified by transcriptional activity. TADs were classified into four quartiles based on the transcriptional activity of genes within each TAD, quantified from nascent RNA-seq signal in control (DMSO) samples, and RAD21 enrichment was averaged across all TADs in each quartile and represented as described in the Methods. RAD21 enrichment correlates with transcriptional activity, with highly transcribed TADs showing increased cohesin retention within TAD interiors and reduced enrichment at TAD boundaries upon SUPT16H depletion, suggesting that transcriptional activity influences cohesin distribution within chromatin domain.

**Figure S5 | Co-depletion of SUPT16H and RAD21 restores transcription in a transcriptional activity-dependent manner. a,** Western blot showing time-course degradation of SUPT16H and RAD21 in RAD21-AID SUPT16H-AID HCT116 cells following treatment with 5-Ph-IAA (1 mM). Both SUPT16H and RAD21 are rapidly depleted within 1 hour of auxin addition. **b,** Metagene analysis of the DMSO/Auxin nascent RNA signal ratio across gene bodies in SUPT16H-AID single degron (red) and SUPT16H-AID RAD21-AID double degron (blue) cells. Genes were first selected based on cohesin accumulation following SUPT16H depletion (RAD21 Auxin/Auxinole ratio >1) and subsequently classified according to transcriptional activity in control (DMSO) samples. Aggregate profiles are shown for all genes, the top 75%, 50%, 25% and 10% most highly transcribed genes in this subset. Lower DMSO/Auxin ratios indicate a smaller transcriptional decrease following auxin treatment. The transcriptional recovery observed upon RAD21 co-depletion becomes more pronounced with increasing transcriptional activity.
